# Warming, but not acidification, increases metabolism and reduces growth of redfish (*Sebastes fasciatus*) in the Gulf of St. Lawrence

**DOI:** 10.1101/2025.01.27.634736

**Authors:** Joëlle J. Guitard, Denis Chabot, Caroline Senay, Dominique Robert, David Deslauriers

**Author notes:** Corresponding author: Joëlle Guitard.

## Abstract

Understanding the effects of global change, including temperature, pH, and oxygen availability, on commercially important species is crucial for anticipating consequences for these resources and their ecosystems. In the Gulf of St. Lawrence (GSL), redfish (Sebastes spp.) have been under moratorium from 1995 to 2024, with a massive recruitment observed in 2011–2013. However, little is known about their metabolic and thermal physiology, making predictions of their response to changing GSL conditions challenging. To address this, we quantified the effects of four acclimatation temperatures (2.5, 5.0, 7.5, and 10.0 ℃) and two pH levels (7.35 and 7.75) on standard and maximum metabolic rates (SMR and MMR), aerobic scope (AS), hypoxia tolerance (O_2_crit), food consumption, and growth in redfish. SMR, MMR, and AS increased with temperature, but growth decreased at the highest temperature, likely due to increased metabolic demand, with food consumption similar across 5.0 to 10.0 °C treatments. O_2_crit was lower for fish acclimated to 2.5 and 5.0 ℃, making redfish less hypoxia-tolerant at higher temperatures. Except from SMR, no significant effect of pH was observed. These results suggest that future changes in the GSL will challenge redfish, with potential long-term effects on their growth due to increased energy requirements.

## Introduction

Global change, through biotic and abiotic factors, impacts marine species with particularly severe effects on boreal and polar species (Parmesan and Yohe 2003; Perry et al. 2005; IPCC 2023). For example, changes in temperature in the North Sea drove a northward change in mean distribution of nearly two-thirds of species (Perry et al. 2005). Such periods of instability can lead to ecosystem regime shifts, which are characterized by drastic changes in food web structure and species composition within the community. One example is the collapse of several pandalid shrimp species under increasing water temperatures in the Gulf of Alaska (Anderson and Piatt 1999), which was accelerated by the resurgence of groundfish species (*Pleuronectidae*, Pacific cod (*Gadus macrocephalus*), Walleye pollock (*Theragra chalcogramma*) as well as pleuronectid flatfish) over the course of a 10-year period (Anderson 2000).

The state of an ecosystem when a major regime shift occurs can greatly influence its resilience (Möllman and Diekmann 2012). While a stable system is typically resilient, one undergoing drastic change in abiotic factors such as temperature or pH is generally more vulnerable to similar changes in species composition and biomass. Georges Bank haddock illustrates how species can recover following a stock collapse (DFO 2024). A 10-fold increase in biomass over 10 years, mostly based on the sole 1999 strong year class (Platt et al. 2003; Head et al. 2005), was observed for this stock, leading to physiological changes, such as reduced growth and a below-average mean size at age (Brodziak et al. 2008). These physiological changes were driven mainly by density-dependent mechanisms and environmental factors like temperature (Wang et al. 2021), and have a major impact on resource availability to the commercial fishery (Brodziak et al. 2008). Understanding species physiological needs in a changing ecosystem through their performance traits helps to better comprehend their impact on their environment.

Performance traits of an individual can determine the ability of an individual to succeed in ecologically relevant tasks such as foraging, competing with other species, growing or reproducing and is defined in this study as the physiological traits investigated (Arnold 1983; Bennett and Huey 1990). In fishery science, growth is a key physiological trait used to assess the health of individuals or cohorts and to predict when a cohort will reach critical size thresholds. These thresholds include the minimum or maximum regulatory size for collection and the length at which 50% of individuals are mature (L50) and considered part of the spawning stock biomass (Brûlé et al. 2024). These indicators are critical for making management decisions such as setting total allowable catch. In addition to growth as a performance trait, aerobic metabolic performance provides valuable insight into the potential energy available to an organism. It has been shown that the level of oxygen uptake is proportional to the amount of energy expenditures in organisms (for reviews, see Chabot et al. 2016; Nelson and Chabot 2024). Aerobic metabolic performance is characterized by an upper and lower bound. The maximal metabolic rate (MMR) represents the maximum oxygen extraction capability (Hulbert and Else, 2000), while the standard metabolic rate (SMR) reflects the minimal energy needed for maintenance in a resting, post-absorptive state (Chabot et al., 2016). The difference between MMR and SMR is known as the aerobic scope (AS), which indicates the maximum energy capacity for activities like locomotion, digestion, growth and reproduction (Claireaux and Lefrançois 2007; Nelson 2016). Consequently, AS can provide insight on the potential energy that an animal can invest towards growth under different environmental constraints when food is not limited.

Most environmental abiotic factors such as temperature (Schulte 2015), dissolved oxygen (Claireaux and Chabot 2016) or pH (Heuer and Grosell 2014) can have a detrimental effect on growth by reducing aerobic scope through multiple pathways for fishes. For the majority of fishes which are ectotherms, temperature acts as a controlling factor which in the case of an increase of temperature accelerates chemical reactions. These changes in temperature can have many effects on different physiological traits of an ectotherm through those processes. For example, it acts differently on standard and maximum metabolic rates and as a result, the relationship between AS and temperature is bell-shaped (Fry 1971; Schulte et al. 2011). The effects of temperature on fish performance traits, such as metabolic rates, growth, and hypoxia tolerance (O_2_crit: the minimum oxygen level required to sustain metabolic demands of maintenance (SMR); Claireaux and Chabot, 2016), are well-documented (Mandic et al. 2009; Gislason et al. 2010; Schulte 2015; Ern et al. 2016). However, performance traits can not only be altered by temperature, but also other abiotic factors such as pH associated with acidification processes. Ocean acidification, caused by elevated CO₂ levels and reduced pH, can act differently on SMR and MMR. Ocean acidification is predicted to act as a masking factor for SMR, meaning that an additional energetic cost will cause a loading stress, which will add to SMR (Heuer and Grosell, 2014). As for MMR, similarly to hypoxia, ocean acidification can act as limiting factor and reduce MMR (Heuer and Grosell, 2014). Given that fish have efficient acid-base regulation mechanisms and substantial variation in sensitivity to acidification, the increase in CO_2_ in the water can, when controlled experimentally, cause contrasting effects in fish (Kreiss et al. 2015; Baumann 2019). For example, in Atlantic cod (*Gadus morhua*) exposed to two temperatures, 18.0 °C and 10.0 °C, an increase in CO_2_ causing acidification induced an increase in SMR for individuals acclimated to 18.0 °C, and a decrease for individuals acclimated at 10.0 °C (Kreiss et al., 2015). In spiny damselfish (*Acanthochromis polyacanthus*) exposed to higher concentrations of CO_2_ (946 µatm CO_2_) lower resting metabolic rates and higher maximal metabolic rates was observed, resulting in a 38% increase of their aerobic scope compared to fish from the control treatment (451 µatm CO_2_) (Rummer et al. 2013). A number of studies have also demonstrated no significant effect of increased CO_2_ on performance traits in adult fish such as aerobic metabolism and growth (for reviews, see Lefevre 2016; Cattano et al. 2018; Yoon et al. 2024). These examples demonstrate that varying levels of CO_2_ in water and changes in temperature can differently impact specific performance traits. Therefore, testing the combined effects of climate change-related abiotic factors is crucial for understanding potential outcomes for fish species.

The GSL is warming at an alarming rate, with the average deep-layer (> 200 m) temperature maximum increasing from 5.2 °C in 2009 to 7.0 °C in 2022 (Galbraith et al. 2024). In highly stratified water bodies such as the GSL, limited oxygen replenishment at depth, combined with high microbial and metabolic activity, accelerates deoxygenation and acidification through the consumption of O₂ and the release of CO₂ by bottom-dwelling organisms (Gilbert et al. 2005; Mucci et al. 2011; Nesbitt and Mucci 2021; Blais et al. 2023). Projections for the 2061-2080 period under the RCP8.5 scenario indicate an increase of about 5 °C in temperature and a decrease of 0.4 on the pH total scale (pH_T_: which accounts for all hydrogen ions, including those in complexes) for the deep-layer of the GSL (Lavoie et al. 2020; IPCC 2023).

Redfish are long-lived, slow-growing, ovoviviparous fish with sporadic recruitment and strong cohorts, that can support commercial fisheries for extended periods (ca. 10 years). Two species, *Sebastes mentella* and *S. fasciatus* are found in the GSL. Visual identification of these species is challenging given their morphological similarities (Senay et al. 2022). One of the last major reproductive events occurred in 1981, supporting fisheries until the early 1990s. However, abnormally cold temperatures in the late 1980s and early 1990s, likely coupled with overfishing, led to recruitment decline and a stock collapse, resulting in the 1995 moratorium. The stock remained low until three successful cohorts emerged in 2011–2013 during a period of rapid warming (Senay et al. 2023). By 2022, redfish represented nearly 82% of the total demersal biomass in Fisheries and Oceans Canada (DFO) bottom trawl surveys, compared to 15% between 1995 and 2012 (Bourdages et al. 2023). This surge not only offers a key fishing opportunity but also impacts the ecosystem through predation and competition (Brown-Vuillemin et al. 2022, 2023). Redfish are also affected by environmental changes, with observed physiological shifts including smaller sizes at maturity (Brûlé et al. 2024) and slower growth rates (Coussau et al. 2024), resulting in redfish attaining the regulatory minimum fishing size of 22 cm at an older age (Senay and Duplisea 2024). Although the actual mechanisms behind these physiological changes remain unclear, factors such as temperature, acidification, reduced dissolved oxygen, and density-dependent mechanisms are likely involved.

Given the close morphological and physiological similarities between *S. fasciatus* and *S. mentella*, *S. fasciatus* serves as a model species to study environmental responses in redfish. This approach is particularly valuable because *S. mentella* typically cannot survive being brought to the surface given that they are found deeper, making comprehensive physiological studies nearly impossible. By using *S. fasciatus* as a model, we aim to infer how both species of redfish might react to changing environmental conditions. Additionally, this research contributes to a broader understanding of how the ecosystem, in which redfish are found, may adapt to global change, providing insights to support improved fishery management and conservation efforts.

Relying on *S. fasciatus* captured in the wild and kept in captivity, the objectives of the present study were to assess the individual and combined effects of projected warming and acidification on performance traits such as growth rate, food consumption, metabolic traits, such as MMR, SMR, AS and O_2_crit in an experimental setting. We were interested in evaluating whether growth, aerobic scope and hypoxia tolerance would decline projected increases in temperature and/or acidification to provide the knowledge necessary to forecast redfish population dynamics in a changing ecosystem.

## Methods

### Fish collection and housing

Redfish were successfully brought live to our aquaculture facilities using the methodology developed by Laroque (2000). Briefly, redfish were collected near *les Escoumins*, Québec, Canada (48.317801°N, 69.413287°W) throughout September and October 2019. SCUBA divers caught ∼16–19 cm redfish with dip nets at a depth of 25–30 m and placed them into cages (52 x 31 x 31 cm), that could hold up to 30 individuals. Once full, cages were placed on the bottom higher up along the slope, at a depth of 10–15 m depending on the tide, to reduce barotrauma. The duration of this equilibration period was at least 12 h, but up to 96 h for fish caught early during each of the 5-day fishing trips. The day after the last capture, the cages were brought to the surface for transportation in oxygenated tanks to the Maurice Lamontagne Institute in Mont-Joli, Québec, Canada. Four fishing trips yielded 822 redfish.

Upon their arrival, fish were transferred into circular water tanks (1 m x 1.1 m, height x diameter; 760 L) supplied with 6 ℃ oxygenated seawater (salinity of 32 ppm). They were offered food (northern shrimp, *Pandalus borealis* and capelin, *Mallotus villosus*) three times a week. It took about two weeks for redfish to acclimatize to their new environment and feed correctly. Wild fish may present various levels of parasite load, which are known to influence performance traits (Chrétien et al. 2022; Guitard et al. 2022). Hence, a week after they started eating, fish received a formaldehyde treatment (37% formaldehyde, 0.01 ml/Liter) for 1 hour to remove possible external parasites. Furthermore, inspection of gills, muscle, and heart when fish were euthanized for tissue collection (n = 32) confirmed the absence of internal parasites.

In December 2019 and January 2020, fish were weighted and individually identified with a PIT-tag (Biomark, Idaho, USA) to allow determination of individual growth. When individuals were anesthetized for length and weight measurements, genetic samples were taken by rubbing a sterile swab for approximately five seconds in their mouth. Each swab was individually placed in a 1.5 ml Eppendorf tube and frozen at −20 °C until species identification was conducted using qPCR, as described in Brûlé et al. (2024). This method accurately assigned individuals to either *S. fasciatus* or *S. mentella* in 96% of cases. Of all individuals included in the experiments, 99.3 % were identified as *S. fasciatus*.

In July 2020, a 2-month experiment compared survival at two salinities, 28 and 32 (50 fish per tank, two tanks per salinity), which resulted in 100% survival in all tanks, therefore salinity was reduced from 32 to 28 (the average salinity of water pumped into the aquaculture facility) over a period of two weeks and then kept at 28 thereafter. After four months of acclimation post salinity experiment, in December 2020, fish were moved to their respective experimental tanks. Experimental methods complied with the regulations of the Canadian Council on Animal Care and were approved by the Maurice Lamontagne Institute animal care committee (Certificate 19-6C). Fish collection in their natural habitat was done under the Canada parks permit (SAGMP-2019-33741).

### Experimental set-up

The experimental set-up was composed of 16 circular water tanks (1 m diameter; 760 L) arranged randomly into 4 temperature lines (2.5, 5.0, 7.5, 10.0 ℃). The four tanks on each line were at the same experimental temperature, with two out of four tanks randomly set to each of the experimental pH levels (7.35 and 7.75). As redfish typically live around 5°C, the temperatures selected included both colder and warmer conditions, with the warmer conditions reflecting the 10°C expected for 2061 from simulation for the GSL by Lavoie et al., 2020. The two pH levels were chosen to represent the actual conditions of the deep layer of the GSL (7.75; Mucci et al. 2011, 2018) and the −0.4 pH units decrease scenario predicted for the 2061-2080 period under an RCP8.5 scenario (7.35; Lavoie et al., 2020, IPCC, 2023). Water temperature was controlled using a custom-built control unit (Gell’Air, Mont-Joli, Canada) with a precision of 0.6 °C in the header tank, but close to 0.1 °C in each of the four tanks within each line. Each line was a semi-open system, fluctuating between 12.5 and 15 liters per minute of new seawater being added to the header tank, and all experimental tanks draining back to the header tank.

A total of 25 fish were randomly distributed in each tank for a total of 400 fish (but see Food Consumption and Growth experiment below for a subsequent change in the number of fish per tank). Fish were fasted for 72 h, anesthetised (Benzocaine, 0.5 ml of solution per liter of seawater, solution = 100 g of benzocaine dissolved in 1 L of 70 % ethanol), weighted, measured (fork length) and then randomly assigned to a tank. Fish were acclimated to the experimental tanks for a week at the same rearing conditions (5 ℃). Then, temperatures were adjusted in increments of 0.5 ℃ per day until all experimental temperatures were reached (*e.g.* 2.5 weeks for 10.0 ℃).

### Monitoring of seawater carbonates

Each tank was equipped with a compression proof glass electrode (model 1208) connected to an Aquastar controller (IKS, Karlsbad, Germany) that allowed for the continuous monitoring and regulation of pH by injecting CO_2_ in the gas exchange column of each tank as required. Additionally, pH on the total scale was verified in each tank daily from Monday to Friday with a pH/conductometer (914, Metrohm AG, Herisau, Switzerland) calibrated once a week using NBS then TRIS (tris(hydroxymethyl)-aminomethane) buffers (to convert pH values to the total scale; Dickson et al. 2007). The same set-up was used for the regulation and monitoring of CO_2_ injection during the respirometry trials: pH on the total scale was verified prior to the respirometry trials, and at least twice during the 4-day respirometry trials.

During the growth experiment, total alkalinity (TA) was monitored every week by sampling water in 104 ml glass bottles in each of the four header tanks (A, B, C, D) for the four different temperature treatments. TA from the header tank was considered the same as that of the four experimental tanks it supplied. TA (in μmol kg^-1^ of seawater) was measured by open-cell potentiometric titration (Mintrop et al. 2000; Dickson 2010) with an automated VINDA sampling system at the Maurice Lamontagne Institute. Each 104 mL sample, warmed to 25 °C, was titrated with 0.1 M hydrochloric acid to the Gran equivalence point (nonlinear curve-fitting method) using a computer-controlled Dosimat (Metrohm AG, USA) dispenser and combination glass electrode. The titrant was calibrated with a series of seawater certified reference materials (CRMs; Andrew Dickson, Scripps Institution of Oceanography, San Diego, CA). The analytical precision of TA was calculated using repeated analyses of duplicate sample and revealed a precision of < 0:1%.

Using the same water samples, pH_T_ was determined weekly in duplicate with an Agilent Cary 60 thermoregulated spectrometer and a 10 cm quartz cell using the purified dye m-cresol purple method (University of South Florida). Absorbance was measured at 730, 578, and 434 nm before and after dye addition at 25 °C (Clayton and Byrne 1993). A similar procedure was conducted before and after each set of sample measurements using a TRIS buffer prepared at a practical salinity (S) value of approximately 30 (Millero 1986). All measurements were converted to the total proton scale (pH_T_) using the measured salinity of each sample and the HSO_4_ association constants given by (Dickson 1990). Reproducibility and accuracy of our measurements were on the order of 0.003 pH units or better.

Both pH_T_ and TA measurements were then used to estimate the aragonite and calcite saturation states, pH_T_ *in situ* and pCO_2_ using the CO_2_SYS program (Lewis and Wallace 1998), and using the dissociation constants (K1 and K2) of Mehrbach et al. (1973), as refit by Dickson and Millero (1987), the KHSO_4_ constant from Dickson (1990), the total boron constant from Lee et al. (2010), and KF constant from Perez and Fraga (1987).

### Food consumption and growth experiment

Fish were fed to satiation three times a week with capelin, northern shrimp, and krill (*Euphausia superba*) each Monday, Wednesday and Friday, respectively. Capelin and shrimp were cut in pieces of approximately 2-3 cm to make them easier to swallow. Food consumption for each tank was measured each feeding day. Food was offered every 5-6 minutes for a minimum of one hour, to ensure that all fish fed *ad libitum*. Portions were weighted before feeding began and uneaten food was collected and used to correct the amount eaten.

Experimental conditions were reached on January 25^th^, 2021, and the end of the experiment was scheduled on June 29^th^, 2021 (5 months), when all fish fasted for a minimum of 72 h, were anesthetized, measured and weighed to calculate growth rate in length and mass. However, slow growth rates and many mass losses were observed between January and June in all treatments, which was not observed in the fish that were not part of the experiment. This was attributed to an initial feeding protocol that did not result in *ad libitum* feeding (feeding stopped as soon as 3 food items accumulated on the bottom in this protocol) and possibly a suboptimal fish density, as redfish are gregarious and stop feeding when fewer than 10 individuals are placed in tanks of this volume (pers. obs.). Thus, on July 13^th^, 17 individuals (afterwards named added fish to differentiate them from the original fish) were added to each experimental tank for a total of 42 fish per tank and 672 fish for the experiment and the feeding protocol was adjusted to ensure *ad libitum* feeding (protocol described at the beginning of this section). Prior to the addition of fish, all experimental tanks were brought to 5 °C over a two to three days depending on the treatment, and injection of CO_2_ in the water was stopped for the 7.35 treatment to increase the pH and help the added fish acclimate to the experimental tanks. Twenty-four hours after the fish addition, temperature and pH were controlled again, and treatment conditions were obtained as of July 16^th^. Day 0 was June 28^th^ for the original fish and July 13^th^ for the added fish and the growth experiment ended on October 20^th^, after 114 and 99 days for both groups, respectively.

Tanks were inspected every day to monitor mortality. The original fish were weighed and measured three times during the whole experiment, on January 25^th^, June 28^th^, and October 20^th^, whereas the added fish were measured twice, on July 14^th^ and October 20^th^.

### Respirometry trials

Two identical separate experimental water baths (200 cm length × 60 cm width × 28 cm height, 336 L) were used, each with an open circulation system. Each water bath was provided with a temperature probe (Hermetically sealed RTD probe, Omega Engineering inc., Norwalk, CT, United States), a controller (1/16 DIN Micromega autotune PID Temperature, Omega Engineering inc.), and two mixing valves (sv3109, Omega Engineering inc.) to enable mixing of the appropriate proportion of cold and warm water (total flow rates between 3.5 and 4.5 L min^-1^) to provide each bath with sea water at the treatment temperature.

Each water bath contained four resting respirometry glass chambers, two of each size (5.214 L; 33 cm, length × 15 cm, diameter or 7.016 L; 49 cm, length × 13.5 cm, diameter). The chambers were transparent but separated with an opaque panel so that fish could not see each other. Each chamber was connected to a closed water circuit to achieve adequate water mixing, made of Tygon tubing and a recirculation pump (model 1048, Eheim, Stuttgart, Germany). The volume of the recirculation loop was 48.5 ml for 5 L chambers and 60 ml for 7 L chambers. Each recirculation loop included housings for a fiber-optic oxygen probe (PyroScience Robust Oxygen) as well as a temperature sensor (PyroScience P100). These were connected to a PyroScience FireSting-O_2_ 4-channel oxygen meter combined with a PyroScience TEX4 (PyroScience GmbH, Aschen, Germany) to have temperature correction on each channel. Dissolved oxygen levels were measured every ∼2 seconds. The four chambers were also connected to a single flush pump (1046, Eheim, Stuttgart, Germany) controlled by the Aquaresp software (V3, Denmark, Aquaresp.com).

At the end of the growth experiment, wet mass-specific oxygen uptake (ṀO_2_, in mg O_2_ h^−1^ kg^−1^) was measured in each respirometer by intermittent-flow respirometry (Steffensen, 1989, Svendsen et al. 2016) using 11.5-minute cycles consisting of a 4-min period of flush to renew the water in the respirometers followed by a 7.5 minute closed period to determine oxygen consumption using the slope of the decreasing dissolved oxygen in the respirometer during the last 6 minutes of the closed phase, when it had become linear.

For each experimental tank, four individuals were chosen randomly among the fish added in July for respirometry trials. Respirometer size was selected as to respect a chamber to fish volume ratio between 20 and 50 (Svendsen et al., 2016) and we obtained a ratio between 29 and 52 with the two respirometer size. We had a total of eight individuals per treatment and 64 fish in total. The respirometry set-up was cleaned between each experiment (every four days) with fresh water and chlorine and was rinsed thoroughly with fresh water and left to run over night with seawater. Oxygen probes were calibrated with 100% and 0% (air sat.) solutions before each experiment. Eight trials were required to cover all eight treatments in duplicate. Treatment tanks were randomly allocated to a given respirometry experiment and water bath.

Prior to all respirometry experiments, fish were fasted for a minimum of 48 hours to ensure they were in a post-absorptive state (Clark et al., 2013; Chabot et al., 2016b). Each trial lasted five days, including preparation and execution. On day one, the set up was cleaned, probes were calibrated (Table S1), fish were selected and weighted and then isolated for the night in a basket floating in the experimental tank. On day two, background oxygen consumption rates (B_*Ṁ*O_2_) were estimated in each empty chamber for a minimum of 40 minutes before and after every respirometry trial. Then, each trial started with a chase protocol followed by 1-minute of air exposure, a common method of estimating MMR in fishes (Roche et al. 2013; Rummer et al. 2016). For this, the selected fish were caught and put in a floating basket in the experimental tank the day prior to the beginning of a respirometry trial. On the day of the trial, each selected fish was individually caught, put in a circular chase area (65 cm × 15 cm, height × diameter, 50 L, depth adjusted as a function of fish height [doubling the height of the fish in water]) using a net and chased by hand until it no longer reacted to contact and tail pinching (chase duration lasted between 1.18 and 8.68 min).The fish was then removed from the arena and held out of the water for 1-minute. The fish was thereafter placed into a respirometry chamber, which was immediately (less than 15 s) sealed. The decline in dissolved oxygen was measured constantly, without flush, until it reached ∼ 80 % air sat. to estimate MMR. Once all four fish had been chased and the MMR estimation completed, control of the system was switched to regular cycles controlled by Aquaresp running the 11.5-minute loops as described above, for the next ∼70 hours (days 3, 4 and 5), with the fish left undisturbed, in the dark (except for a daily check-up with a red light), behind opaque curtains. Over this period, oxygen uptake rates of fish stabilized, and SMR could be estimated. Oxygen levels remained above 95% in the chambers during all cycles of all trials.

After ∼60 hours (day 5), oxygen uptake under decreasing dissolved oxygen (DO) was quantified to estimate hypoxia tolerance (O_2_crit). This was achieved by gradually injecting nitrogen in the respirometry bath. DO was lowered at a rate of approximately 5 % between each cycle until DO in the chamber reached approximately 30 % sat. Then DO was lowered at a rate of 2-3 % between cycles. Aquaresp, paired with an R script, calculated MO_2_ in real time and the experiment lasted until the animal could not sustain SMR estimated during the previous days (Claireaux and Chabot, 2016). We visually checked on the animals to make sure they were not in distress or lost equilibrium during this part of the experiment. Hypoxia tolerance trials lasted between 7 and 9 hours. This protocol follows best practices for collecting and reporting respirometry data as described in Killen et al. (2021) (see Table S1).

### Data extraction and analysis

#### Food consumption and Growth rates

All data were analysed using R v. 4.2.0 (R Foundation for Statistical Computing, 2022). Food consumption was analysed as follows from July 13^th^ to October 20^th^:

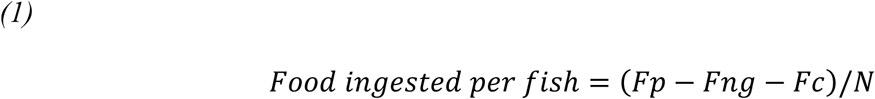

where *Food ingested per fish* is the amount of food in grams eaten by one individual of a given tank for one meal (capelin, shrimp or krill), *Fp* is the amount of food prepared for that tank for that meal, *Fng* is the amount of food not given, *Fc* is the amount of food collected at the bottom of the tank at the end of the meal and *N* is the number of fish in the tank during that meal.

Food consumption was calculated as the amount ingested by one fish per week. Since redfish consistently ate more capelin and less shrimp, it was preferable to calculate weekly food consumption as the sum of per capita consumption for the three meals:

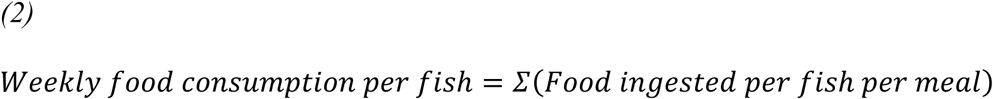

Growth rates were calculated on all fish present at the end of the experiment (October 20^th^). Specific growth rate (Jobling, 1983) was calculated for each individual from June 28^th^ for the original fish and from July 13^th^ for the added fish to October 20^th^.

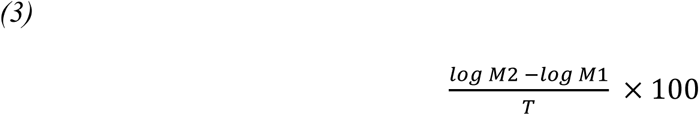

where M2 is the mass at the end of the experiment (October 20^th^), M1 the mass at the beginning of the experiment (June 28^th^ or July 14^th^) and T the duration of the experiment (114 days for the original fish, 99 days for the added fish).

#### Respirometry data

Wet mass-specific oxygen uptake (*Ṁ*O_2_; mg O2 h^-1^ kg^-1^) were calculated from the slopes of the linear regressions between oxygen concentration and time, accounting for the volume of water in the respirometer (respirometer gross volume less fish mass as an estimate of fish volume, assuming a density of 1g ml^-1^) for the 64 individuals tested.

Following the chase and air exposure protocol, oxygen uptake was measured during approximately 30 min without any flush/reoxygenation period because oxygen uptake changes quickly after the chase and the maximum rate could otherwise occur during a flush. To further reduce the risk of underestimating MMR, rolling regressions were used to detect the steepest part of the long decline in DO in the respirometer (Zhang et al. 2019). Contrary to sequential regressions, when one interval begins after the end of the previous interval, in rolling regressions each interval begins one point after the beginning of the previous interval. The shortest reliable interval or window width (WW) over which these regressions should be calculated was determined according to the methodology outlined by Zhang et al. (2019) and modified by Guscelli et al. (2023). This analysis gives the shortest reliable WW of 3 min (See section on MMR determination in Supplemental information).

SMR was estimated with the *fishMO2* package (Chabot 2020) as the quantile (p = 0.2) of the *Ṁ*O_2_ values obtained after an acclimatation period of 16 h and the start of the decline in DO used to measure O_2_crit (Chabot et al. 2016b). Because we have observed night activity in this species, we have estimated SMR for each individual with and without including nights. We then have selected the lowest estimation of SMR out of these two rates. The background respiration rate (B_*Ṁ*O_2_) was subtracted from the *Ṁ*O_2_ measurements assuming a linear increase in bacterial respiration from the start to the end of the trial.

Absolute AS was calculated as the difference between MMR and SMR (Fry 1948; 1971). O_2_crit was identified with data points measured below SMR. A linear regression was fitted to the oxygen uptake values below SMR, at low values of DO, and the intersection of this line and the horizontal line representing SMR was taken to be O_2crit_ (Claireaux and Chabot 2016; Chabot 2020).

#### Statistical analysis

To test the effect of temperature (average temperature during the growth experiment) and pH on performance traits such as food consumption and growth, and because these variables had non-linear relationships, we used a full Generalized Additive Model (GAM) framework with the mgcv package (Wood 2017).

Food intake was measured for whole tanks, and it was not possible to distinguish the food intake of original and added fish. The full model included pH as a fixed factor and one smooth for each pH level, in addition to experimental tank as a random factor (Tables S3, Model Food1). Food intake models and their AIC are shown in Table S3, with description of all terms in Table S2. Model selection was based on the Akaike Information Criterion (AIC) and the model with the lowest AIC was the final model.

In the model elaboration for growth rates, as the original group of 25 fish had been subjected to experimental conditions for a longer period compared to the fish added on July 13th, group (original and added) was included as a fixed factor in the growth model. The first model tested was the full model, which included pH, group and their interaction as fixed factors and allowed for a different smooth for growth rate as a function of temperature foreach combination of pH and group. Experimental tank was included as a random factor. The interaction and the group term were progressively removed (Table S3). Selection of the final model proceeded as for food consumption models.

Preliminary models for oxygen uptake were formulated with the gam() function, but the relationship between SMR, MMR or AS and temperature were always linear. For this reason, linear mixed effects models (lmer function in lme4 package; Bates et al. 2015; Table S3) were used for the three metabolic traits (MMR, SMR, AS) as well as hypoxia tolerance (O2crit) where temperature (average temperature during each respirometry experiment) was a continuous variable, pH was a 2-level factor and the experimental tank and respirometry chamber were two random variables. The full models included the interaction between temperature and pH as a test of homogeneity of slopes, an assumption of ANCOVAs (Quinn and Keough 2002). Reduced models without interaction were calculated when the P-value of the interaction was not significant. R-squared values for the lme models were determined using the MuMIn package (Barton 2024). Two fish were rejected from the models on respirometry data for two different reasons. One individual escaped from the chamber during the first night and was immediately reintroduced in the chamber the following morning. Unfortunately, this event caused added stress, and this individual did not reach SMR during the remainder of the session. Another fish was discarded because its oxygen uptake was still declining at the end of the period used to estimate SMR (Fig. S4).

Model assumptions for all models were assessed visually using diagnostic plots with the DHARMa package (Hartig 2022) and were met for all models.

## Results

### Survival rate

Survival rate for the experimental period ranged from 90 to 100 % (death of 0 to 4 individuals per tank; Table S3; S4) for the whole experiment. The 13 deaths did not appear related to the treatments (Table S4) and in fact were either accidental (fish that jumped out through imperfections in the net covering the tank) or euthanasia of fish that developed exophthalmia, to respect animal care procedures.

### Growth performance

#### Food consumption

The final simplified GAM for food consumption included pH as a fixed effect, with a smoothed effect of temperature (mean temperature throughout the experiment) that varied by pH and experimental tank was included as a random effect (Table S3; Model Food 2).

Food consumption ranged from 0.81 g to 13.27 g fish^-1^ week^-1^ (mean 7.21 g fish^-1^ week^-1^, SD 2.42) across all tanks. Food consumption was not influenced by pH as a categorical variable (p = 0.215; Table 1) however, it was lower at 2.5 °C than at the three higher temperatures (p = 0.004; Table 1; Fig. 1A).

**Figure 1.**
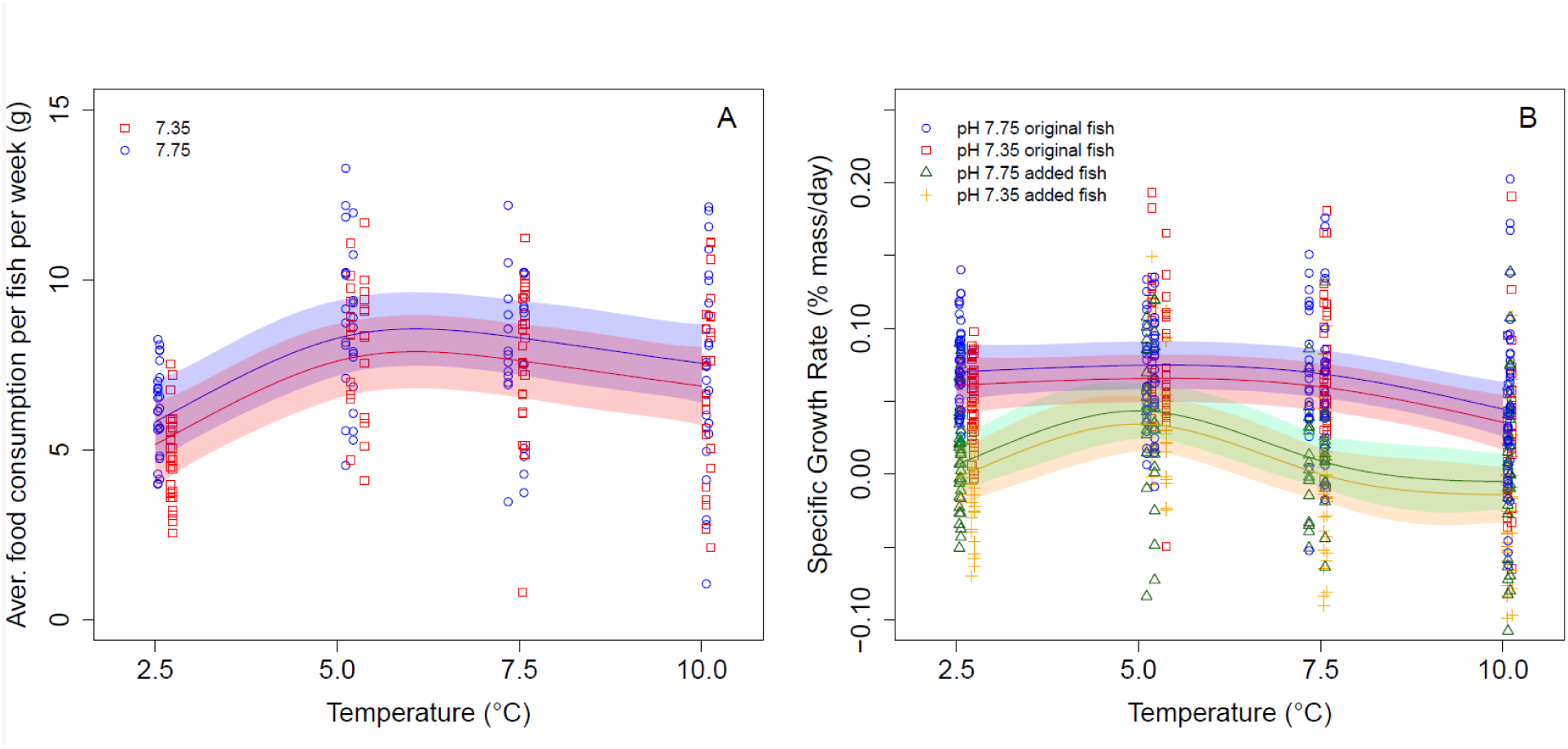
Effect of temperature and pH on food consumption and growth rates. **(A)** Food consumption per fish in grams per day for redfish (*S. f*asciatus) reared at temperature between 2.5 °C and 10.0 °C in seawater with current GSL pH (blue) and seawater reduced by 0.4 units using CO_2_ (red). N= 208 **(B)** Specific growth rates in redfish after acclimation to temperature between 2.5 °C and 10.0 °C in seawater with current GSL pH (blue and green) and seawater reduced by 0.4 units using CO_2_ (red and yellow) N=662. **The shaded areas around each regression lines represents the 95% confidence intervals.**

**Table 1.** Relationship between food consumption or growth and temperature and pH, with tank as random factor. Test statistics obtained from generalized additive model (GAM) of growth rates and food consumption as a function of temperature, pH, and group with tank treatment as a random factor for redfish. Statistically significant results are indicated in bold. edf: estimated degrees of freedom, F-value: F-statistic for the smooth term, p-value: statistical significance threshold. Statistically significant results are indicated in bold with significance threshold of 0 ‘***’ 0.001 ‘**’ 0.01 ‘*’ 0.05 ‘.’ 0.1 ‘ ’ 1.

| Response | Predictors | edf | f-value | p-value |
| --- | --- | --- | --- | --- |
| **Growth rates** |  |  |  |  |
| (g day<sup>-1</sup>) | pH |  | 1.108 | 0.293 |
|  | Group |  | 231.153 | **<0.0001\*\*\*** |
|  | Temp*added fish | 2.894 | 6.569 | **<0.0001\*\*\*** |
|  | Temp*original fish | 2.026 | 3.665 | **0.020\*** |
|  | Tank | 10.243 | 5.266 | **<0.0001\*\*\*** |
| **Food consumption** |  |  |  |  |
| (g week<sup>-1</sup>) | pH |  | 1.301 | 0.215 |
|  | Temperature | 2.149 | 3.636 | **0.004\*\*** |
|  | Tank | 9.309 | 9.667 | **<0.0001\*\*\*** |

#### Specific growth rates in mass

The final simplified GAM for specific growth rates included pH and group as fixed effects and mean temperature throughout the experiment by group as smoothed fixed effects (degree of smoothing k=9), experimental tank was included as a random effect (Table S3; Model Growth 5).

Across all treatments, fish had a mean mass on beginning of the experiment (June 28^th^ for the original fish and July 13^th^ for the added fish) of 158.42 g (SD = 38.84 g) for the original and 188.05 g (SD = 53.82 g) for the added fish, with an overall (added and original) mean mass of 170.74 g (SD = 47.91 g). Across all treatments, growth rates ranged from −0.06 to 0.20 % body mass day^-1^ with a mean of 0.06 % body mass day^-1^ (SD = 0.04 % body mass day^-1^) for the original fish and ranged from −0.129 to 0.149 % body mass day^-1^ with a mean of 0.009 % body mass day^-1^ (SD = 0.051 % body mass day^-1^) for the added fish. Growth was not influenced by pH (p =0.293, Table 1), but it was influenced by the group (p < 0.0001; Table 1) and the temperature smooth term for each group was significant (original fish: p = 0.020, added fish: p = 0.0001; Table 1). Further, the shape of the relationship between growth rate and temperature differed for each group (Fig. 1B). Deviance explained for this model was 40.6%. Specific growth rate was highest at 5.0 °C and lowest at 10.0 °C for both groups (Fig. 1B). pH did not influence growth rate. For the original fish, which were acclimated to the treatments before the growth experiment, growth rates were significantly lower at 10.0 °C compared to the three lower temperature treatments. Added fish, which were subjected to the treatments at the beginning of the growth trial, had lower growth rates at 2.5 °C and 10.0 °C, peaking around 5.0 °C.

### Aerobic metabolic performance

#### MMR, SMR, AS

The final simplified LME model for metabolic traits tested the effect of temperature and pH as fixed effects and included the respirometry chamber and the experimental tank, from the growth experiment, as random effects (Table S3; Model SMR2-MMR2-AS2-O_2_crit2).

Mean mass of individuals before respirometry trials was 176.77 g (SD = 34.97). MMR ranged from 104 to 216 mg O_2_ h^-1^ kg^-1^ while SMR ranged from 15 to 51 mg O_2_ h^-1^ kg^-1^. MMR and AS were not influenced by pH (p_MMR_ = 0.8517; p_AS_ = 0.6102; Fig. 2; Table 2), however SMR was significantly elevated for the acidic pH (7.35) at all temperatures (p_pH_= 0.0257, p_Temp_ < 0.0001, Fig. 2B, Table 2). There was a significant positive relationship between temperature and the three metabolic traits (Fig. 2; Table 2), fish exposed to higher temperatures had higher MMR, SMR and AS. The random effect “tank” added to the model was statistically significant (p ≤ 0.05) for MMR, AS and O_2_crit, meaning that the experimental tank in which fish were reared during the growth experiment influenced those traits.

**Figure 2.**
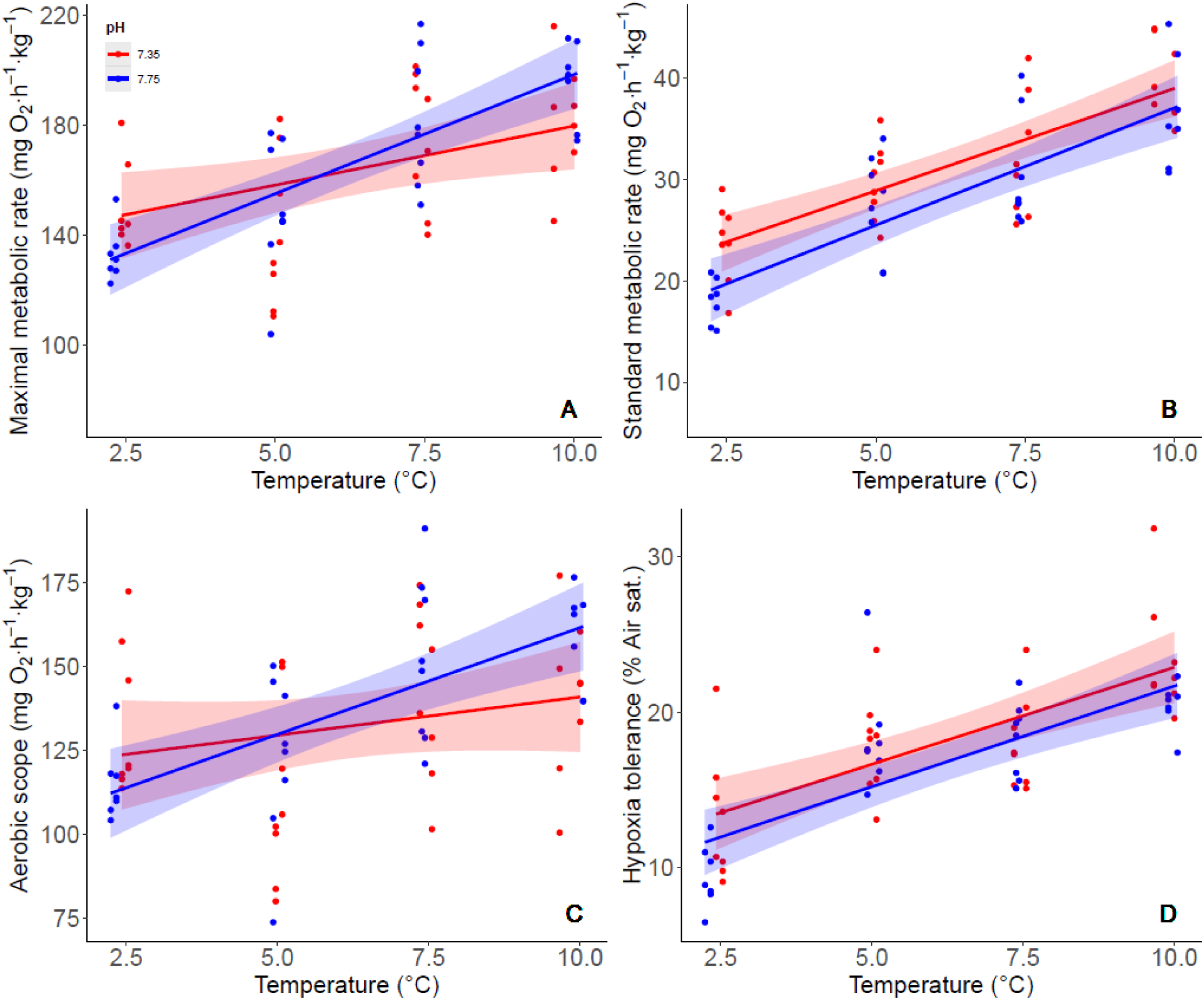
Effect of temperature and pH on metabolic traits. Oxygen uptake rates (mg O2 h^-1^ kg^-1^) as a function of exposition temperature and pH (red acidic: 7.35 and blue control: 7.75) in redfish (*S. fasciatus*). Maximal metabolic rate **(A)**, standard metabolic rate **(B)**, aerobic scope **(C)** and hypoxia tolerance **(D)**. **The shaded areas around each regression lines represents the 95% confidence intervals. The slopes for the two pH levels did not differ significantly in A–D.**

**Table 2.** Relationship between metabolic traits estimated, temperature, pH with the tank and the respirometry chamber as random factors. Test statistics obtained from linear mixed-effects model for maximum metabolic rate (MMR), standard metabolic rate (SMR), aerobic scope (AS) and hypoxia tolerance (O_2_crit) as a function of temperature and pH treatment in redfish (n=62). Statistically significant results are indicated in bold with significance threshold of 0 ‘***’ 0.001 ‘**’ 0.01 ‘*’ 0.05 ‘.’ 0.1 ‘ ’ 1. Marginal R^2^ (R^2^M) and conditional R^2^ (R^2^C) indicate the variance explained by fixed factors and by both fixed and random factors, respectively (Nakagawa and Schielzeth, 2013).

| Response | Predictors | Chisq | DF | p-value | R<sup>2</sup>M | R<sup>2</sup>C |
| --- | --- | --- | --- | --- | --- | --- |
| **MMR** |  |  |  |  | 0.35 | 0.54 |
| (mgO<sub>2</sub>h<sup>-1</sup>kg<sup>-1</sup>) | Temperature | 20.687 | 1 | **<0.0001\*\*\*** |  |  |
|  | pH | 0.0349 | 1 | 0.8517 |  |  |
| **SMR** |  |  |  |  | 0.64 | 0.69 |
| (mgO<sub>2</sub>h<sup>-1</sup>kg<sup>-1</sup>) | Temperature | 77.831 | 1 | **<0.0001\*\*\*** |  |  |
|  | pH | 4.972 | 1 | **0.0257\*** |  |  |
| **AS** |  |  |  |  | 0.18 | 0.47 |
| (mgO<sub>2</sub>h<sup>-1</sup>kg<sup>-1</sup>) | Temperature | 8.1683 | 1 | **0.0042\*\*** |  |  |
|  | pH | 0.2598 | 1 | 0.6102 |  |  |
| **O<sub>2</sub>crit** |  |  |  |  | 0.51 | 0.63 |
| (% Air sat.) | Temperature | 37.2667 | 1 | **<0.0001\*\*\*** |  |  |
|  | pH | 1.5734 | 1 | 0.2097 |  |  |

#### Hypoxia tolerance (O_2_crit)

Hypoxia tolerance for the temperature range of this experiment was between 6.5 % air sat (2.5 ℃) and 31.8 % air sat (10.0 ℃) (Fig. 2D). There was no relationship between O_2_crit and pH. There was a significant positive relationship between O_2_crit and temperature (p < 0.0001, Fig. 2D, Table 2). Individuals acclimated at 2.5 ℃ and 10.0 ℃ became unable to maintain SMR at ca. 14 % and 22 % air sat, respectively, a difference of 8 % air saturation between the extreme temperatures.

## Discussion

To the best of our knowledge this is the first study to conduct experimental work on redfish, given the significant challenge of bringing them to the laboratory alive and in good health. Our results showed that the predicted deep-layer temperature for the GSL in 2061 (10.0 °C) affected redfish through food consumption, growth and metabolic traits, although we did not observe an effect of acidic pH on most of the performance traits of redfish. In this context, growth was decreased at the highest temperature (10 °C) likely due to an increase in metabolic demand, which could not be sustained by sufficient food consumption even though individuals were fed *ad libitum*.

### Energy allocation

In fish bioenergetics, when modelling energy allocation, consumed energy is first allocated to metabolism (maintenance, activity, digestion), then to waste (feces and urine) and the left over can be allocated to somatic or gonadal growth (Madenjian et al. 2023). As temperature increases, it is common to observe an increase in aerobic metabolism until the organism reaches a thermal threshold. Before this threshold is reached, higher temperatures can cause some functions such as postprandial increase in oxygen uptake (Specific Dynamic Action: SDA), to require more of that energy (Chabot et al. 2016a; Nelson and Chabot 2024). Consequently, other functions, such as growth, become lower energetic priorities. In recent years, Jutlfet et al. (2021) proposed the *aerobic scope protection* hypothesis This hypothesis suggests that fish consuming the same size meal experience a higher SDA peak in warm water, which intensifies at even higher temperatures, potentially reducing the aerobic scope available for other oxygen-demanding activities. As Jutfelt et al. (2021) stated, animals should then ideally regulate their food intake based on their need to maintain aerobic scope under challenging conditions that temperature increase can cause. As a result, warming temperature should result in decreased food intake as well as reduced growth, if food intake decreases while aerobic metabolism remains high. In the present experiment, SDA was not estimated for redfish under the different temperature treatments. However, the stability of food consumption at the highest two temperature treatments, when metabolic demands increased, suggested that such a process might be occurring.

Two secondary results emerged from our study. First, we have observed that for growth the two groups (original fish and added) responded similarly to the highest temperature (10℃) but had different trends for lower temperatures. The major difference is that added fish had a lower growth rate at 2.5 ℃ in comparison with the 5.0 ℃ treatment while the original fish had similar growth rates for these two temperatures. This could be explained by the fact that the added fish had less time to acclimate to the experimental conditions compared to the original fish that were kept in these conditions for four months prior to data collection. This acclimation time difference also led to overall significantly lower growth rates for the added fish (Table 1, Fig. 1), but the temperature-growth rate relationship showed similar conclusions for both groups when it came to predicting the effects of warmer waters expected for the second half of the Century in this region.

Second, even though growth rates were generally reduced at 10 ℃, one of the four experimental tanks exposed to that temperature showed one of the best mean growth rates of all 16 tanks, causing a large variability at this temperature. This tank was also the one out of the four 10 ℃ treatment that had the highest food consumption (Fig. S5). The hypothesis behind the Aerobic scope protection stated by Jutfelt et al. (2021) relates to behaviour, where fish would willingly regulate their food intake based on the additional energetic costs associated to an increase in temperature. However, given that we did not observe a decline in aerobic scope at 10 ℃, our results suggest that this tank might not have adopted this strategy, and instead simply ate more than the others resulting in faster growth rate. We thus speculate that for fish kept at 10°C, those able to consume more food can grow faster. However, this is considering that individuals are reared in regulated tanks without any stressors such as predation risk, etc and that such observations might be different for fish in the wild. The lowest food consumption rates throughout the experiment were observed at either 10°C for the three other tanks or 2.5°C (Fig. S5), indicating that, at similar food quantities, temperature plays a significant role in regulating growth, particularly when comparing the two extreme treatments.

### Food consumption and growth

The highest acclimation temperature in our experiment (10.0 ℃) led to lower growth rates. We observed that fish maintained relatively constant food consumption at temperatures ranging from 5.0 to 10.0 ℃, while experiencing increased metabolic demands. Consequently, this suggests that even when fed *ad libitum*, a reduction in growth is to be anticipated at high temperatures in the wild. Moreover, in natural environments characterized by density-dependent pressures, it is highly probable that the effect of high temperature would be magnified under limited prey supply (Sánchez Lizaso et al. 2000). The current reduction in growth measured in GSL redfish in the wild (Senay et al., 2023, Coussau et al., 2024) could be partly attributable to density dependence under increasing temperatures, caused by the sharp decline of the Northern shrimp (Bourdages et al., 2022), recognized as a preferred prey (Brown-Vuillemin et al. 2022; 2023). Interestingly, the lowest food consumption in this study happened at the lowest temperature tested (2.5 ℃) but this did not result in the slowest growth rate, which can be explained by the lower metabolic demand. This suggests that GSL redfish is well adapted to a cold, subarctic environment and that a lower food consumption rate induced by density-dependence would not be as aggravating at low temperatures (2.5℃) relative to those predicted for the GSL (10.0℃).

### Aerobic metabolism

We observed that higher acclimation temperatures within our study range resulted in higher MMR, SMR and AS. Out of the hypotheses that have proposed a mechanistic understanding for the relationship between aerobic scope and growth, the oxygen- and capacity-limited thermal tolerance (OCLTT) theory is a widely -recognized framework. The OCLTT speculates that as an animal approaches its thermal limits, its ability to supply oxygen to tissues decreases, leading to a gradual decline in aerobic performance and eventually reaching lethal temperatures (Pörtner, 2010). This occurs because higher temperatures drive a continuous increase in oxygen demand, which the animal’s oxygen supply system can no longer meet. This theory also states that aquatic ectotherms have evolved biochemical and physiological adaptations to maximize aerobic metabolism within a specific temperature range, thereby optimizing fitness-related performance such as growth and reproduction (Pörtner 2001). Consequently, from the OCLTT point of view the temperature at which the optimum aerobic scope is observed will also be the optimal temperature for growth (Pörtner 2010). Our results do not support this hypothesis and are more in line with that proposed by Clark et al. (2013) and Schulte et al. (2011) who demonstrated that when tested experimentally, many species’ aerobic scopes and other physiological traits do not align with the thermal profiles of other critical traits like growth and reproduction (e.g. Gräns et al. 2014; Norin et al. 2014). In our study, an increase in aerobic scope did not translate into increased growth for the same temperatures suggesting that the concept of optimal temperature varies among physiological traits (Fig. 1 and 2).

As we observed a linear increase in MMR, SMR, and AS with temperature, the acclimation temperatures used in our study did not capture the maximum thermal range for redfish aerobic metabolism. However, we are confident that the thermal limit was not exceeded, which would have resulted in a decline in these metabolic traits at higher temperatures. Temperature selection in the present study was based on knowledge of redfish distribution in the GSL, where they are systematically found at temperatures below 8 °C. We thus hypothesized that 10 °C would be sufficiently high to observe a decline in metabolic performance and a more pronounced reduction in growth rate. Although *S. fasciatus* appears to prefer water temperatures around 5-6 °C in the wild where they are found in higher densities (Kelly et al. 1972; Atkinson 1989; Senay et al. 2023), they have also been found in warmer waters and up to 13 °C in the Gulf of Maine (Scott, 1982). Future studies should expose *S. fasciatus* to higher temperatures to better define the optimum temperature range for metabolic rates and growth.

### Acidification

The effects of ocean acidification, estimated through a decrease in pH, on aerobic metabolic traits have been documented but have shown contrasting results in fishes (Heuer and Grosell 2014; Esbaugh 2018), including in the genus Sebastes. Copper rockfish (*S. caurinus*) that were exposed for several weeks to two pH levels (normal: 7.87 vs low: 7.32) were characterized by a reduction in their critical aerobic swimming speed (16.8%) and aerobic scope (53.5%) under low pH comparatively with normal pH. However, no changes were observed in these traits for blue rockfish (*S. mystinus*) exposed to the same conditions and coming from the same habitat (Hamilton et al. 2017). Even though these two species are closely related and use the same habitat as adults, this difference could be explained by the fact that blue rockfish were exposed to high pCO_2_ levels during early life stages, possibly leading to a better adaptation for these conditions than copper rockfish (Hamilton et al. 2017). Given that redfish migrate through the water column in earlier life stages, one hypothesis is that they might have been exposed to different pH levels as the water column of the deep water of the GSL is well stratified and surface layers tend to be more acidic (Lavoie et al. 2021). This exposition of redfish larvae to lower pH levels in the surface layer may result in better tolerance to low pH later in life.

As Heuer & Grosell (2014) stated, the main effect of water acidification on aerobic metabolism, when observed, is primarily that of a loading factor, *sensu* Fry (1971): when acidification had an effect, SMR increased (cost of acid-base regulation), driving a reduction in AS and subsequently, growth. In this study, we did observe that fish acclimated at an acidic pH had a higher SMR than those at the control pH, fitting this hypothesis. Therefore, in the context of ocean acidification, redfish may potentially see their AS and their growth potential decreased. Probably because of the considerable variation in SMR, MMR and AS in this study, the increase in SMR did not have any measurable impact on aerobic scope, even when combined with an increase in temperature. In a similar study on Atlantic halibut, Gräns et al. (2014) observed elevated aerobic scope at lower pH (acidic) but no effect on growth rates except at their lower temperature treatment. At this lower temperature treatment (5 °C) growth rate was suppressed by 24% at the low pH compared to current pH. Surprisingly, this lower temperature treatment is the temperature where these juveniles halibut are typically found in the wild (Gräns et al., 2014).

### Hypoxia tolerance

Hypoxia tolerance, measured as O_2_crit, is the ambient oxygen level at which a fish can meet its maintenance requirements but has no aerobic scope left for activity (Claireaux and Chabot, 2019). In the present study, fish acclimated to higher temperatures had lower hypoxia tolerance, reflected in a higher O_2_crit. As oxygen levels continue to drop, the fish is dependent on anaerobic metabolism, which is unsustainable in time. Rapid decreases in dissolved oxygen as those simulated in the laboratory are unlikely to happen in the redfish habitat, but this measurement indicates the oxygen limit that redfish can face in their environment. As redfish are a competitor to other groundfish in the GSL, O_2_crit makes it possible to compare the oxygen sensitivity of redfish to that of competitors such as Atlantic cod (*Gadus morhua*) or Greenland halibut (*Reinhardtius hippoglossoides*). Plante et al. (1998), working with Gulf of St. Lawrence cod of various sizes exposed to 2 and 6 °C, observed complete mortality within hours at 10% air saturation and found that only a few individuals survived 96 hours at 16% air saturation, highlighting the species’ sensitivity to hypoxia. Adult Greenland halibut kept at 5 °C were able to tolerate ambient dissolved oxygen down to 11% air sat., though juveniles were more sensitive to hypoxia (Dupont-Prinet et al. 2013). During our experiments we observed that redfish had a mean O_2_crit of 18.59 % at 5.0 °C and 22.64 % at 10.0 °C, making them more tolerant at higher temperatures than cod, but more sensitive to hypoxia than Greenland halibut at water temperatures where all three species are frequently observed. Considering that dissolved oxygen levels currently range between 15 and 65% sat. in the deep channels (Blais et al. 2023; Wallace et al. 2023), some areas, particularly the estuary, are already suboptimal habitat for redfish. Worsening of hypoxia is observed in the deep channels of the GSL and estuary (Gilbert et al. 2005; Jutras et al. 2020), and predicted to continue (Lavoie et al., 2020), likely to cause even more habitat loss for GSL redfish, especially as this will be accompanied by warming, which makes redfish more sensitive to hypoxia (this study).

### Perspectives and conclusion

The results of this study are timely as the fishery for GSL redfish has just reopened in 2024, which highlights the importance of a better understanding of redfish physiological responses under multiple environmental constraints. The GSL will continue to be influenced by anthropogenic inputs in multiple ways, causing warming, acidification, and low oxygen levels. Linking the growth patterns and food requirements for growth with a mechanistical approach should allow us to better predict the future of this species in its ecological role in the ecosystem and as a key resource for fisheries.

The potential redistribution of redfish within the water column, driven by warming temperatures and reduced oxygen levels, could alter their ecological role in the GSL. Adult redfish currently distribute at the bottom of the deep layer of the GSL, where waters are warming the most. The GSL waters are stratified, with a colder cold intermediate layer (CIL) located between the deep and surface layers. This stratification implies the possibility that redfish move up in the water column to access more favorable temperature for growth near the CIL. Such vertical redistribution could have broad implications to cold-associated species which would undergo increased predation risk from redfish.

From a fisheries perspective, anticipated future slower growth and smaller fish sizes present two challenges: a reduction in biomass, which could affect catch volume, and potential reduction in the value of the catch as smaller fish are worth less economically than larger fish and are therefore resulting in lower landing value. Although reduced fecundity has not been directly observed, limited food availability coupled with higher metabolic demands may constrain energy allocation, potentially affecting both growth and reproduction. This could hamper future recruitment, with implications for the long-term sustainability of the population and the viability of the fishery.

Future work should investigate the species’ responses to higher temperature thresholds, variable prey availability, prolonged exposure to reduced oxygen levels but also SDA under these different climate change scenarios. Developing a mechanistic understanding of these processes, particularly the impact of digestion on aerobic scope and growth, will be essential for predicting how redfish populations may adapt to the rapidly changing conditions of the GSL. This knowledge will inform conservation efforts and management strategies aimed at sustaining both the ecological and economic value of redfish in the region.

## Author contributions

JG: Conceptualization, Data curation, Formal analysis, Funding acquisition, Investigation, Methodology, Project administration, Software, Validation, Visualization, Writing - original draft; DC: Conceptualization, Data curation, Formal analysis, Funding acquisition, Methodology, Project administration, Resources, Software, Supervision, Validation, Writing - review & editing; CS: Funding acquisition, Writing - review & editing; DR: Conceptualization, Funding acquisition, Supervision, Validation, Writing - review & editing; DD: Conceptualization, Funding acquisition, Methodology, Project administration, Resources, Supervision, Validation, Writing - review & editing.

## Acknowledgements

The authors wish to thank D. Picard, T. Hansen, F. Hartog, K. MacGregor, JD. Tourangeau-Larivière and J. Heinerth for technical support during the experiment at the DFO Maurice Lamontagne Institute (Mont-Joli, QC, Canada) and for fish collection. The authors wish to acknowledge the Indigenous Peoples and the history of the traditional territories on which our work was conducted. J.G., D.D. and D.R. are members of the Ressources Aquatiques Québec inter-institutional strategic research network.

## Data availability

Data generated or analyzed during this study are available in the Université du Québec à Rimouski’s Dataverse Collection within the Borealis Repository. DOI to come

## References

Anderson, P.J. 2000. Pandalid shrimp as indicators of ecosystem regime shift. J. Northwest Atl. Fish. Sci. 27(1880): 1–10. doi:10.2960/j.v27.a1.

Anderson, P.J., and Piatt, J.F. 1999. Community reorganization in the Gulf of Alaska following ocean climate regime shift. Mar. Ecol. Prog. Ser. 189: 117–123. doi:10.3354/meps189117.

Arnold, S.J. 1983. Morphology, performance and fitness. Am. Zool. 361: 347–361.

Atkinson, D.B. 1989. Seasonal distribution of sharp-beaked redfish (Sebastes spp.) in northeast Grand Bank. J. Northwest Atl. Fish. Sci. 9(2): 141–150. doi:10.2960/J.v9.a13.

Barton, K. 2024. _MuMIn: Multi-Model Inference_. R package version 1.48.4, <https://CRAN.R-project.org/package=MuMIn>.

Bates, D., Mächler, M., Bolker, B., and Walker, S. 2015. Fitting linear mixed-effects models using lme4. J. Stat. Softw. 67(1): 1–48. 10.18637/jss.v067.i01.

Baumann, H. 2019. Experimental assessments of marine species sensitivities to ocean acidification and co-stressors: How far have we come?1. Can. J. Zool. 97(5): 399–408. doi:10.1139/cjz-2018-0198.

Bennett, A.F., and Huey, R.B. 1990. Studying the evolution of physiological performance. *In* In: Futuyma DJ, Antonovics J, Oxford., Oxford sur. *Edited by* D. Futuyma and J. Antonovics. Oxford, UK, University Press. p. 251.

Blais, M., Galbraith, P.S., Plourde, S., and Lehoux, C. 2023. Chemical and Biological Oceanographic Conditions in the Estuary and Gulf of St. Lawrence during 2022. Can. Tech. Rep. Hydrogr. Ocean Sci. 357: v +70 p.

Bourdages, H., Brassard, C., Desgagnés, M., Galbraith, P.S., Gauthier, J., Nozères, C., Scallon-Chouinard, P.-M., and Senay, C. 2023. Preliminary results from the ecosystemic survey in August 2022 in the Estuary and northern Gulf of St. Lawrence. In DFO Canadian Science Advisory Secretariat Research Document 2023/074 iv +100p.

Brodziak, J., Traver, M.L., and Col, L.A. 2008. The nascent recovery of the Georges Bank haddock stock. Fish. Res. 94: 123–132. doi:10.1016/j.fishres.2008.03.009.

Brown-vuillemin, S., Chabot, D., Nozères, C., Sirois, P., and Robert, D. 2022. Diet composition of red fish (Sebastes sp.) during periods of population collapse and massive resurgence in the Gulf of St. Lawrence. Front. Mar. Sci.: 1–21. doi:10.3389/fmars.2022.963039.

Brown-vuillemin, S., Robert, D., Tremblay, R., Chabot, D., and Sirois, P. 2023. Feeding ecology of redfish (Sebastes sp.) inferred from the integrated use of fatty acid profiles as complementary dietary tracers to stomach content analysis. J. Fish Biol. 102: 1049–1066. doi:10.1111/jfb.15348.

Brûlé, C., Benhalima, K., Roux, M.-J., Parent, G., Chavarria, C., and Senay, C. 2024. Updating knowledge of Redfish (Sebastes mentella and S. fasciatus) reproduction in a changing environment, the Gulf of St. Lawrence and Laurentian Channel, Canada. J. Fish Biol. (January): 1–20. doi:10.1111/jfb.15677.

Cattano, C., Claudet, J., Domenici, P., and Milazzo, M. 2018. Living in a high CO 2 world: a global meta-analysis shows multiple trait-mediated fi sh responses to ocean acidi fi cation. Ecol. Monogr. 88(3): 320–335. doi:10.1002/ecm.1297.

Chabot, D. 2020. fishMO2: Calculate and plot the standard metabolic rate (SMR), the critical oxygen level (O2crit) and the specific dynamic action (SDA) and related variables in fishes and crustaceans, v. 0.46. https://rdrr.io/github/denis-chabot/fishMO2/man/fishMO2-pack.

Chabot, D., Koenker, R., and Farrell, A.P. 2016a. The measurement of specific dynamic action in fishes. J. Fish Biol. 88(1): 152–172. doi:10.1111/jfb.12836.

Chabot, D., Steffensen, J.F., and Farrell, A.P. 2016b. The determination of standard metabolic rate in fishes. J. Fish Biol. 88(1): 81–121. doi:10.1111/jfb.12845.

Chrétien, E., De Bonville, J., Guitard, J., Binning, S.A., Melis, É., Kack, A., Côté, A., Gradito, M., Papillon, A., Thelamon, V., Levet, M., and Barou-Dagues, M. 2022. Few studies of wild animal performance account for parasite infections: A systematic review. J. Anim. Ecol. (July): 1–13. doi:10.1111/1365-2656.13864.

Claireaux, G., and Chabot, D. 2016. Responses by fishes to environmental hypoxia: Integration through Fry’s concept of aerobic metabolic scope. J. Fish Biol. 88(1): 232–251. doi:10.1111/jfb.12833.

Claireaux, G., and Lefrançois, C. 2007. Linking environmental variability and fish performance: Integration through the concept of scope for activity. Philos. Trans. R. Soc. B Biol. Sci. 362(1487): 2031–2041. doi:10.1098/rstb.2007.2099.

Clark, T.D., Sandblom, E., and Jutfelt, F. 2013. Aerobic scope measurements of fishes in an era of climate change: respirometry, relevance and recommendations. J. Exp. Biol. 216(15): 2771–2782. doi:10.1242/jeb.084251.

Clayton, D.H., and Byrne, R.H. 1993. Spectrophotometric seawater pH measurements: total hydrogen ion concentration scale calibration of m-cresol purple and at-sea results. Deep. Res. Part I Oceanogr. Res. Pap. 40: 2115–2129.

Coussau, L., Morissette, O., Robert, D., and Sirois, P. 2024. Drivers of growth in strong year classes of the deepwater redfish (Sebastes mentella) population from the Gulf of St. Lawrence derived from otolith increment-based growth chronologies. J. Fish Biol. (July). doi:10.1111/jfb.15903.

DFO. 2024. Eastern Georges Bank haddock (Melanogrammus aeglefinus) assessment to 2023.

Dickson, A.G. 1990. Thermodynamics of the dissociation of boric acid in synthetic seawater from 273.15 to 318.15 K. Deep. Res. Part I Oceanogr. Res. Pap. 37: 755–766.

Dickson, A.G. 2010. The carbon dioxide system in seawater: Equilibrium chemistry and measurements. In Guide to best practices for ocean acidification research and data reporting. Edited by U. Riebesell, V.J. Fabry, and L. Hansson. Publications Office of the European Union, Luxembourg. pp. 17–52.

Dupont-Prinet, A., Vagner, M., Chabot, D., and Audet, C. 2013. Impact of hypoxia on the metabolism of Greenland halibut (Reinhardtius hippoglossoides). Can. J. Fish. Aquat. Sci. 70(3): 461–469. doi:10.1139/cjfas-2012-0327.

Ern, R., Norin, T., Gamperl, A.K., and Esbaugh, A.J. 2016. Oxygen dependence of upper thermal limits in fishes. J. Exp. Biol. 219(21): 3376–3383. doi:10.1242/jeb.143495.

Esbaugh, A.J. 2018. Physiological implications of ocean acidification for marine fish: emerging patterns and new insights. J. Comp. Physiol. B Biochem. Syst. Environ. Physiol. 188(1): 1–13. Springer Berlin Heidelberg. doi:10.1007/s00360-017-1105-6.

Fry, F.E.J. 1971. The effect of environmental factors on the physiology of fish. Fish Physiol. 6(C): 1–98. doi:10.1016/S1546-5098(08)60146-6.

Galbraith, P.S., Chassé, J., Shaw, J., Dumas, J., and Bourassa, M.-N. 2024. Physical Oceanographic Conditions in the Gulf of St. Lawrence during 2023. Can. Tech. Rep. Hydrogr. Ocean Sci. 378 : v +91 p.

Gilbert, D., Sundby, B., Gobeil, C., Mucci, A., and Tremblay, G.H. 2005. A seventy-two-year record of diminishing deep-water oxygen in the St. Lawrence estuary: The northwest Atlantic connection. Limnol. Oceanogr. 50(5): 1654–1666. doi:10.4319/lo.2005.50.5.1654.

Gislason, H., Daan, N., Rice, J.C., and Pope, J.G. 2010. Size, growth, temperature and the natural mortality of marine fish. Fish Fish. 11(2): 149–158. doi:10.1111/j.1467-2979.2009.00350.x.

Gräns, A., Jutfelt, F., Sandblom, E., Jönsson, E., Wiklander, K., Seth, H., Olsson, C., Dupont, S., Ortega-Martinez, O., Einarsdottir, I., Björnsson, B.T., Sundell, K., and Axelsson, M. 2014. Aerobic scope fails to explain the detrimental effects on growth resulting from warming and elevated CO2 in Atlantic halibut. J. Exp. Biol. 217(5): 711–717. doi:10.1242/jeb.096743.

Guitard, J.J., Chrétien, E., De Bonville, J., Roche, D.G., Boisclair, D., and Binning, S.A. 2022. Increased parasite load is associated with reduced metabolic rates and escape responsiveness in pumpkinseed sunfish. J. Exp. Biol. 225(15): 1–12. doi:10.1242/jeb.243160.

Guscelli, E., Noisette, F., Chabot, D., Blier, P.U., Hansen, T., Ros, M.C., Pepin, P., Skanes, K.R., and Calosi, P. 2023. Northern shrimp from multiple origins show similar sensitivity to global change drivers, but different cellular energetic capacity. J. Exp. Biol. doi:10.1242/jeb.245400.

Hamilton, S.L., Logan, C.A., Fennie, H.W., Sogard, S.M., Barry, J.P., Makukhov, A.D., Tobosa, L.R., Boyer, K., Lovera, C.F., and Bernardi, G. 2017. Species-specific responses of juvenile rockfish to elevated pCO2: From behavior to genomics. PLoS One 12(1): 1–23. doi:10.1371/journal.pone.0169670.

Hartig, F. 2022. _DHARMa: Residual Diagnostics for Hierarchical (Multi-Level / Mixed) Regression Models_. R package version 0.4.6, <https://CRAN.R-project.org/package=DHARMa>.

Head, E.J.H., Brickman, D., and Harris, L.R. 2005. An exceptional haddock year class and unusual environmental conditions on the Scotian Shelf in 1999. J. Plankton Res. 27(6): 597–602. doi:10.1093/plankt/fbi019.

Heuer, R.M., and Grosell, M. 2014. Physiological impacts of elevated carbon dioxide and ocean acidification on fish. Am. J. Physiol. - Regul. Integr. Comp. Physiol. 307(9): R1061–R1084. doi:10.1152/ajpregu.00064.2014.

IPCC. 2023. Sections. In: Climate Change 2023: Synthesis Report. Contribution of Working Groups I, II and III to the Sixth Assessment Report of the Intergovernmental Panel on Climate Change [Core Writing Team, H. Lee and J. Romero (eds.)]. IPCC, Geneva, Switzerland,.

Jutfelt, F., Norin, T., Åsheim, E., Rowsey, L., Andreassen, A., Morgan, R., Clark, T., and Speers-Roesch, B. 2021. Aerobic scope protection reduces ectotherm growth under warming. Funct. Ecol. (April): 1–11. doi:10.1111/1365-2435.13811.

Kelly, G.F., Earl, P.M., and Kaylor, J.D. 1972. Redfish. In NOAA NMFS Ext. Fish. Facts Publ. No. 1.

Kreiss, C.M., Michael, K., Lucassen, M., Jutfelt, F., Motyka, R., Dupont, S., and Pörtner, H.O. 2015. Ocean warming and acidification modulate energy budget and gill ion regulatory mechanisms in Atlantic cod (Gadus morhua). J. Comp. Physiol. B 185(7): 767–781. Springer Berlin Heidelberg. doi:10.1007/s00360-015-0923-7.

Laroque, R. 2000. A SCUBA technique for collecting live Sebastes spp. specimens. Can. Tech. Rep. Fish. Aquat. Sci. 2309 v +13 p.

Lavoie, D., Lambert, N., Rousseau, S., Dumas, J., Chassé, J., Long, Z., Perrie, W., Starr, M., Brickman, D., and Azetsu-Scott, K. 2020. Projections of future physical and biogeochemical conditions in the Gulf of St. Lawrence, on the Scotian Shelf and in the Gulf of Maine. In Can. Tech. Rep. Hydrogr. Ocean Sci.

Lavoie, D., Lambert, N., Starr, M., Chassé, J., Riche, O., Le Clainche, Y., Azetsu-Scott, K., Béjaoui, B., Christian, J.R., and Gilbert, D. 2021. The Gulf of St. Lawrence Biogeochemical Model: A Modelling Tool for Fisheries and Ocean Management. Front. Mar. Sci. 8(October): 1–29. doi:10.3389/fmars.2021.732269.

Lee, K., Kim, T., Byrne, R.H., Millero, F.J., Feely, R.A., and Liu, Y. 2010. The universal ratio of boron to chlorinity for the North Pacific and North Atlantic oceans. Geochim. Cosmochim. Acta 74(6): 1801–1811. Elsevier Ltd. doi:10.1016/j.gca.2009.12.027.

Lefevre, S. 2016. Are global warming and ocean acidification conspiring against marine ectotherms? A meta-analysis of the respiratory effects of elevated temperature, high CO2 and their interaction. Conserv. Physiol. 4(1): 1–31. doi:10.1093/conphys/cow009.

Lewis, E., and Wallace, D.W.R. 1998. Program Developed for CO2 System Calculations. arbon Dioxide Inf. Anal. Center, Oak Ridge Natl. Lab. U.S. Dep. Energy, Oak Ridge, TN, USA: 42.

Madenjian, C.P., Chipps, S.R., Deslauriers, D., Guitard, J.J., and Daigle, N.J. 2023. Fish bioenergetics modeling. In Reference Module in Life Sciences, Second Edi. Elsevier. doi:10.1016/b978-0-323-90801-6.00063-x.

Mandic, M., Todgham, A.E., and Richards, J.G. 2009. Mechanisms and evolution of hypoxia tolerance in fish. Proc. R. Soc. B Biol. Sci. 276(1657): 735–744. doi:10.1098/rspb.2008.1235.

Mehrbach, C., Culberson, C.H., Hawley, J.E., and Pytkowicx, R.M. 1973. Measurement of the apparent dissociation constants of carbonic acid in seawater at atmospheric pressure. Limnol. Oceanogr. 18(November): 897–907.

Millero, F.J. 1986. The pH of estuarine waters. Limnol. Oceanogr. 31(4): 839–847.

Mintrop, L., Pérez, F.F., González-Dávila, M., Santana-Casiano, M., and Körtzinger, A. 2000. Alkalinity determination by potentiometry: Intercalibration using three different methods. Ciencias Mar. 26: 23–27. 10.7773/cm.v26i1.573.

Möllman, C., and Diekmann, R. 2012. Marine Ecosystem Regime Shifts Induced by Climate and Overfishing : A Review for the Northern Hemisphere. In Advances in ecological research, Global change in multispecies systems: Part II. Edited by G. Woodward, U. Jacob, and E.J. O’Gorman. Academic Press, Burlington. pp. 303–347. doi:10.1016/B978-0-12-398315-2.00004-1.

Mucci, A., Starr, M., Gilbert, D., and Sundby, B. 2011. Acidification of Lower St. Lawrence estuary bottom waters. Atmos. - Ocean 49(3): 206–218. doi:10.1080/07055900.2011.599265.

Nelson, J.A. 2016. Oxygen consumption rate v. rate of energy utilization of fishes: A comparison and brief history of the two measurements. J. Fish Biol. 88(1): 10–25. doi:10.1111/jfb.12824.

Nelson, J.A., and Chabot, D. 2024. Energy consumption : Metabolism. In Encyclopedia of Fish Physiology from mechanism to application, Second Edition. Edited by S.L. Alderman and T.E. Gillis. Elsevier, Academic Press. pp. 482–492.

Nesbitt, W.A., and Mucci, A. 2021. Direct evidence of sediment carbonate dissolution in response to bottom-water acidification in the Gulf of St. Lawrence, Canada. Can. J. Earth Sci. 58(1): 84–92. doi:10.1139/cjes-2020-0020.

Norin, T., Malte, H., and Clark, T.D. 2014. Aerobic scope does not predict the performance of a tropical eurythermal fish at elevated temperatures. J. Exp. Biol. 217(2): 244–251. doi:10.1242/jeb.089755.

Parmesan, C., and Yohe, G. 2003. A globally coherent fingerprint of climate change. Nature 421: 37–42.

Perez, F.F., and Fraga, F. 1987. Association constant of fluoride and hydrogen ions in seawater. Mar. Chem. 21(July 1987): 161–168. doi:10.1016/0304-4203(87)90036-3.

Perry, A.L., Low, P.J., Ellis, J.R., and Reynolds, J.D. 2005. Climate change and distribution shifts in marine fishes. Science (80-.). 308: 1912–1915. doi:10.1126/science.1111322.

Platt, T., Fuentes-Yaco, C., and Frank, K.T. 2003. Spring algal bloom and larval fish survival. Nature 423: 398–399. doi:10.1016/j.jaci.2012.05.050.

Pörtner, H. 2001. Climate change and temperature-dependent biogeography: Oxygen limitation of thermal tolerance in animals. Naturwissenschaften 88(4): 137–146. doi:10.1007/s001140100216.

Pörtner, H.O. 2010. Oxygen- And capacity-limitation of thermal tolerance: A matrix for integrating climate-related stressor effects in marine ecosystems. J. Exp. Biol. 213(6): 881–893. doi:10.1242/jeb.037523.

Quinn, G.P., and Keough, M.J. 2002. Experimental design and data analysis for biologists. Cambridge university press, Cambridge.

Rummer, J.L., Stecyk, J.A.W., Couturier, C.S., Watson, S.A., Nilsson, G.E., and Munday, P.L. 2013. Elevated CO2 enhances aerobic scope of a coral reef fish. Conserv. Physiol. 1(1): 1–7. doi:10.1093/conphys/cot023.

Sánchez Lizaso, J.L., Goñi, R., Reñones, O., García Charton, J.A., Galzin, R., Bayle, J.T., Sánchez Jerez, P., Perez Ruzafa, A., and Ramos, A.A. 2000. Density dependence in marine protected populations: A review. Environ. Conserv. 27(2): 144–158. doi:10.1017/S0376892900000187.

Schulte, P.M. 2015. The effects of temperature on aerobic metabolism: Towards a mechanistic understanding of the responses of ectotherms to a changing environment. J. Exp. Biol. 218(12): 1856–1866. doi:10.1242/jeb.118851.

Schulte, P.M., Healy, T.M., and Fangue, N.A. 2011. Thermal Performance Curves, Phenotypic Plasticity, and the Time Scales of Temperature Exposure. Integr. Comp. Biol. 51(5): 691–702. doi:10.1093/icb/icr097.

Senay, C., Bermingham, T., Parent, G.J., Hugues, P., Parent, É., and Bourret, A. 2022. Identifying two Redfish species, Sebastes mentella and S. fasciatus, in fishery and survey catches using anal fin ray count in Units 1 and 2. Can. Tech. Rep. Fish. Aquat. Sci. 3445 viii +:46.

Senay, C., and Duplisea, D. 2024. Potential Removals of Redfish (Sebastes mentella and S. fasciatus) in Unit 1. DFO Can. Sci. Advis. Secr. Res. Doc. 2024/031 iv+:11.

Senay, C., Rousseau, S., Brûlé, C., Chavarria, C., Isabel, L., Parent, G.J., Chabot, D., and Duplisea, D. 2023. Unit 1 Redfish (Sebastes mentella and S. fasciatus) stock status in 2021. DFO Can. Sci. Advis. Sec. Res. Doc. 2023/036. xi +125. Available from https://publications.gc.ca/collections/collection_2023/mpo-dfo/fs70-5/Fs70-5-2023-036-eng.pdf.

Wallace, D.W.R., Jutras, M., Nesbitt, W.A., and Donaldson, A. 2023. Can green hydrogen production be used to mitigate ocean deoxygenationA scenario from the Gulf of St. Lawrence. Mitig. Adapt. Strateg. Glob. Chang. 28(8): 1–15. Springer Netherlands. doi:10.1007/s11027-023-10094-1.

Wang, Y., Gharouni, A., Friedland, K.D., and Melrose, D.C. 2021. Effect of environmental factors and density-dependence on somatic growth of Eastern Georges Bank haddock (Melanogrammus aeglefinus). Fish. Res. 240(August 2020): 105954. Elsevier B.V. doi:10.1016/j.fishres.2021.105954.

Wood, S.. 2017. Generalized Additive Models: An Introduction with R (2nd edition). Chapman and Hall/CRC.

Yoon, G.R., Bozai, A., and Porteus, C.S. 2024. Could future ocean acidification be affecting the energy budgets of marine fish ? Conserv. Physiol. 12: 1–13. doi:10.1093/conphys/coae069.

Zhang, Y., Gilbert, M.J.H., and Farrell, A.P. 2019. Finding the peak of dynamic oxygen uptake during fatiguing exercise in fish. J. Exp. Biol. 222(November). doi:10.1242/jeb.196568.

