## Supplemental information for "Warming, but not acidification, increases metabolism and reduces growth of redfish (*Sebastes fasciatus*) in the Gulf of St. Lawrence"

**Table S1. Checklist of 53 essential criteria for the reporting of methods for aquatic intermittent-flow respirometry**.

| Number | Criterion and Category | Response | Value (where required) | Units |
|---|---|---|---|---|
| | **EQUIPMENT, MATERIALS, AND SETUP** | | | |
| **1** | Body mass of animals at time of respirometry | Yes | | |
| **2** | Volume of empty respirometers | Yes | 5 and 7 liters | |
| **3** | How chamber mixing was achieved | Yes, with a pump through recirculation loop | | |
| **4** | Ratio of net respirometer volume (plus any associated tubing in mixing circuit) to animal body mass | Yes chamber/fish | | |
| **5** | Material of tubing used in any mixing circuit | Yes, Tygon | | |
| **6** | Volume of tubing in any mixing circuit | Yes | 48.5 ml for 5 L and 60 ml for 7 L | |
| **7** | Confirm volume of tubing in any mixing circuit was included in calculations of oxygen uptake | YES | | |
| **8** | Material of respirometer (e.g. glass, acrylic, etc.) | Yes, glass | | |
| **9** | Type of oxygen probe and data recording | Yes | | |
| **10** | Sampling frequency of water dissolved oxygen | Yes | 2 | seconds |
| **11** | Describe placement of oxygen probe (in mixing circuit or directly in chamber) | Yes, mixing circuit | | |
| **12** | Flow rate during flushing and recirculation, or confirm that chamber returned to normoxia during flushing | Yes | | |
| **13** | Timing of flush/closed cycles | Yes. | | |

| | | | | |
|---|---|---|---|---|
| **14** | Wait (delay) time excluded from closed measurement cycles | Yes | | |
| **15** | Frequency and method of probe calibration (for both 0 and 100% calibrations) | Yes | Once every trial | Sodium sulphite was used to obtain a 0% solution for calibration. Bubblers were used to calibrate for 100%, Calibrations were done before each trial at the temperature of the trial. |
| **16** | State whether software temperature compensation was used during recording of water oxygen concentration | | | |
| | **MEASUREMENT CONDITIONS** | | | |
| **17** | Temperature during respirometry | Yes and included in model | | |
| **18** | How temperature was controlled | Yes | | |
| **19** | Photoperiod during respirometry | Yes. No photoperiod. Behind black curtains throughout | | |
| **20** | If (and how) ambient water bath was cleaned and aerated during measurement of oxygen uptake (e.g. filtration, periodic or continuous water changes) | Yes | Once between every trial | |
| **21** | Total volume of ambient water bath and any associated reservoirs | Yes | 336 | Liters |
| **22** | Minimum water oxygen dissolved oxygen reached during closed phases | yes | 80 | % air sat |
| **23** | State whether chambers were visually shielded from external disturbance | Yes | | |
| **24** | How many animals were measured during a given respirometry trial (i.e. how many animals were in the same water bath) | Yes | 4 in the same water bath, 2 | |

| | | | | |
|---|---|---|---|---|
| | | | separated bath, total of 8 | |
| **25** | If multiple animals were measured simultaneously, state whether they were able to see each other during measurements | Yes. | They could not see each other. Panel separating them | |
| **26** | Duration of animal fasting before placement in respirometer | yes | 72 | Hours |
| **27** | Duration of all trials combined (number of days to measure all animals in the study) | Yes | 8 trials of 5 days | |
| **28** | Acclimation time to the laboratory (or time since capture for field studies) before respirometry measurements | Yes | See: Growth experiment | |
| | **BACKGROUND RESPIRATION** | | | |
| **29** | State whether background microbial respiration was measured and accounted for, and if so, method used (e.g. parallel measures with empty respirometry chamber, measurements before and after for all chambers while empty, both) | Yes | Before and after trials | |
| **30** | State if background respiration was measured at beginning and/or end, state how many slopes and for what duration | Yes | Beginning and end | 5 slopes minimum on each end |
| **31** | State how changes in background respiration were modelled over time (e.g. linear, exponential, parallel measures) | Yes | linear | |
| **32** | Level of background respiration (e.g. as a percentage of SMR) | | | |
| **33** | Method and frequency of system cleaning (e.g. system bleached between each trial, UV lamp) | Yes, specified in methods | | |
| | **STANDARD OR ROUTINE METABOLIC RATE** | | | |
| **34** | Acclimation time after transfer to chamber, or alternatively, time to reach beginning of metabolic rate measurements after introduction to chamber | Yes | 16 | Hours |
| **35** | Time period, within a trial, over which oxygen uptake was measured (e.g. number of hours) | Yes | 3 ½ | days |
| **36** | Value taken as SMR/RMR (e.g. quantile, mean of lowest 10 percent, mean of all values) | Yes | 0.2 quantile (Chabot et al., 2016) | |
| **37** | Total number of slopes measured and used to derive metabolic rate (e.g. how much data were used to calculate quantiles) | Between 0 and 10% of slopes were rejected during the | | |

| | | | | |
|---|---|---|---|---|
| | | period estimated for SMR | | |
| **38** | Whether any time periods were removed from calculations of SMR/RMR (e.g. data during acclimation, periods of high activity [e.g. daytime]) | Due to some nocturnal spontaneous activity, nights were excluded for 44 individuals | | |
| **39** | $r^2$ threshold for slopes used for SMR/RMR (or mean) | 0.93 in one case but mostly 0.95 | | |
| **40** | Proportion of data removed due to being outliers below r-squared threshold | No outliers | | |
| | **MAXIMUM METABOLIC RATE** | | | |
| **41** | When MMR was measured in relation to SMR (i.e. before or after) | Yes, before | | |
| **42** | Method used (e.g. critical swimming speed respirometry, swim to exhaustion in swim tunnel, or chase to exhaustion) | Yes, chase + air | | |
| **43** | Value taken as MMR (e.g. the highest rate of oxygen uptake value after transfer, average of highest values) | Yes, highest rate of oxygen uptake over the trial | | |
| **44** | If MMR measured post-exhaustion, length of activity challenge or chase (e.g. 2 min, until exhaustion, etc.) | Yes | | |
| **45** | If MMR measured post-exhaustion, state whether further air-exposure was added after exercise | Yes | 1 | minute |
| **46** | If MMR measured post-exhaustion, time until transfer to chamber after exhaustion or time to start of oxygen uptake recording | Yes | | |
| **47** | Duration of slopes used to calculate MMR (e.g. 1 min, 5 min, etc.) | Yes | | |
| **48** | Slope estimation method for MMR (e.g. rolling regression, sequential discrete time frames) | Yes. Rolling regression | | |
| **49** | How absolute aerobic scope and/or factorial aerobic scope is calculated (i.e. using raw SMR and MMR, allometrically mass-adjusted SMR and MMR, or allometrically mass-adjusting aerobic scope itself) | Yes | | |
| | **DATA HANDLING AND STATISTICS** | | | |
| **50** | Sample size | **Yes** | 64 | |

| | | |
|---|---|---|
| **51** | How oxygen uptake rates were calculated (software or script, equation, units, etc.) | Yes |
| **52** | Confirm that volume (mass) of animal was subtracted from respirometer volume when calculating oxygen uptake rates | Yes |
| **53** | State whether analyses accounted for variation in body mass and describe any allometric mass-corrections or adjustments | Yes |

**Table S2. Temperature and pH variation in the experimental tanks from February 22nd to July 1st, 2021.**

| **Treatment** T °C_pH | **Tank** | **Mean temperature (°C)** | **Standard error** | **Mean pH** | **Standard error** |
|---|---|---|---|---|---|
| **5.0_7.35** | **A1** | 5.18 | 0.03 | 7.34 | 0.01 |
| **5.0_7.75** | **A2** | 5.07 | 0.03 | 7.73 | 0.00 |
| **5.0_7.75** | **A3** | 4.97 | 0.03 | 7.72 | 0.00 |
| **5.0_7.35** | **A4** | 5.04 | 0.03 | 7.34 | 0.01 |
| **7.5_7.75** | **B1** | 7.53 | 0.03 | 7.73 | 0.00 |
| **7.5_7.35** | **B2** | 7.53 | 0.03 | 7.33 | 0.00 |
| **7.5_7.35** | **B3** | 7.49 | 0.03 | 7.35 | 0.01 |
| **7.5_7.75** | **B4** | 7.37 | 0.03 | 7.72 | 0.00 |
| **2.5_7.35** | **C1** | 2.59 | 0.02 | 7.34 | 0.01 |
| **2.5_7.35** | **C2** | 2.54 | 0.02 | 7.34 | 0.01 |
| **2.5_7.75** | **C3** | 2.45 | 0.03 | 7.73 | 0.00 |
| **2.5_7.75** | **C4** | 2.43 | 0.02 | 7.73 | 0.00 |
| **10.0_7.75** | **D1** | 10.13 | 0.02 | 7.72 | 0.00 |
| **10.0_7.35** | **D2** | 10.15 | 0.02 | 7.35 | 0.00 |
| **10.0_7.75** | **D3** | 10.10 | 0.02 | 7.72 | 0.00 |
| **10.0_7.35** | **D4** | 10.09 | 0.02 | 7.35 | 0.00 |

**Table S3. Temperature and pH variation in the experimental tanks from July 2nd to October 20th, 2021, and respirometry during trial.**

| **Treatment** T °C_pH | **Tank** | **Mean temperature experiment (°C)** | **Standard error** | **Mean pH experiment** | **Standard error** | **Mean temperature respirometry session (°C)** | **Standard error** | **Mean pH respirometry session** | **Standard error** |
|---|---|---|---|---|---|---|---|---|---|
| **5.0_7.35** | **A1** | 5.61 | 0.04 | 7.36 | 0.01 | 4.85 | 0.00 | 7.38 | 0.00 |
| **5.0_7.75** | **A2** | 5.43 | 0.04 | 7.67 | 0.00 | 5.09 | 0.00 | 7.73 | 0.01 |
| **5.0_7.75** | **A3** | 5.33 | 0.04 | 7.67 | 0.00 | 5.07 | 0.00 | 7.76 | 0.01 |
| **5.0_7.35** | **A4** | 5.41 | 0.03 | 7.37 | 0.01 | 5.11 | 0.00 | 7.35 | 0.04 |
| **7.5_7.75** | **B1** | 7.57 | 0.03 | 7.71 | 0.00 | 7.30 | 0.00 | 7.85 | 0.09 |
| **7.5_7.35** | **B2** | 7.59 | 0.02 | 7.37 | 0.01 | 7.53 | 0.00 | 7.38 | 0.02 |
| **7.5_7.35** | **B3** | 7.6 | 0.02 | 7.35 | 0.00 | 7.69 | 0.00 | 7.36 | 0.02 |
| **7.5_7.75** | **B4** | 7.33 | 0.03 | 7.72 | 0.00 | 7.39 | 0.00 | 7.76 | 0.01 |
| **2.5_7.35** | **C1** | 2.85 | 0.03 | 7.36 | 0.01 | 2.43 | 0.00 | 7.37 | 0.01 |
| **2.5_7.35** | **C2** | 2.83 | 0.03 | 7.36 | 0.01 | 2.41 | 0.00 | 7.35 | 0.03 |
| **2.5_7.75** | **C3** | 2.59 | 0.03 | 7.71 | 0.01 | 2.38 | 0.00 | 7.72 | 0.02 |
| **2.5_7.75** | **C4** | 2.58 | 0.03 | 7.71 | 0.01 | 2.35 | 0.01 | 7.78 | 0.00 |
| **10.0_7.75** | **D1** | 10.07 | 0.04 | 7.69 | 0.00 | 10.08 | 0.00 | 7.75 | 0.01 |
| **10.0_7.35** | **D2** | 10.12 | 0.04 | 7.36 | 0.00 | 10.07 | 0.00 | 7.33 | 0.02 |
| **10.0_7.75** | **D3** | 10.05 | 0.04 | 9.95 | 0.00 | 7.73 | 0.00 | 7.69 | 0.00 |
| **10.0_7.35** | **D4** | 10.08 | 0.04 | 9.92 | 0.00 | 7.31 | 0.01 | 7.36 | 0.00 |

**Table S3. Description and variable type of all parameters used in the generalized additive models for each physiological trait**

| Parameter | Description | Variable type |
|---|---|---|
| **Growth** | | |
| Temperature | Mean temperature during the whole growth experiment as a continous variable | Fixed |
| pH | pH treatment as a 2 level factor (7.35 or 7.75) | Fixed |
| Group | Group at which fish belong to during the growth experiment. Original or added | Fixed |
| pH*Group | Interaction between pH and Group | Fixed |
| pH.Group | pH treatment for each group as a 4 level factor (original 7.35, original 7.75, added 7.35, added 7.35) | Fixed |
| Tank | Tank in which fish spent the whole growth experiment | Random |
| **Food consumption** | | |
| Temperature | Mean temperature during the whole growth experiment as a continous variable | Fixed |
| pH | pH treatment as a 2 level factor (7.35 or 7.75) | Fixed |
| Tank | | Random |
| **Metabolic traits** | | |
| **(MMR, SMR, AS, $O_{2crit}$)** | | |
| Temperature | Mean temperature during the respirometry trial for each treatment as a continous variable | Fixed |
| pH | pH treatment as a 2 level factor (7.35 or 7.75) | Fixed |
| Temperature*pH | Interaction between temperature and pH | Fixed |
| Tank | Tank in which fish spent the whole growth experiment | Random |
| Respirometry chamber | Respirometry chamber in which the fish underwent respirometry as a 8 level factor | Random |

**Table S4. Design for best generalized additive models and linear mixed-effects models for each physiological trait with AIC. Model selection was based on the lowest AIC value and choosing the most parsimonious model when the interaction was not significant.**

| **Model** | **AIC** | **P-Value** |
|---|---|---|
| Growth 1 = gam(Growth rates ~ pH*group + s(Temperature, k=9, by=interaction(pH, group, drop=TRUE)) + s(tank, bs="re", data= data, method="REML") | -2283.817 | |
| Growth 2 = gam(Growth rates ~ pH + group + s(Temperature, k=9, by=interaction(pH, group, drop=TRUE)) +s(tank, bs="re"), data= data, method="REML") | -2285.064 | |
| Growth 3 = gam(Growth rates ~ pH + s(Temperature, k=9, by=interaction(pH, group, drop=TRUE)) + s(tank, bs="re"), data= data, method="REML") | -2258.626 | |
| Growth 4 = gam(Growth rates ~ pH + group + s(Temperature, k=9, by=pH) + s(tank, bs="re"), data= data, method="REML") | -2281.605 | |
| Growth 5 = gam(Growth rates ~ pH + group + s(Temperature, k=9, by=group) + s(tank, bs="re"), data= data, method="REML") | -2292.213 | |
| Growth 6 = gam(Growth rates ~ pH + group + s(Temperature, k=9) + s(tank, bs="re"), data= data, method="REML") | -2283.131 | |
| Food 1 = gam(Weekly food consumption~pH+s(Temperature, k=9, by=pH) + s(tank, bs="re"), data=data, method="REML") | 899.6192 | |
| Food 2 = gam(Weekly food consumption~pH+s(Temperature, k=9) + s(tank, bs="re"), data=data, method="REML") | 897.6241 | |
| SMR 1 = lmer(SMR ~ Temperature*pH + (1 \| tank) + (1\|respirometer), data=data) | | 0.556 |
| SMR 2 = lmer(SMR ~ Temperature + pH + (1 \| tank) + (1\|respirometer), data=data) | | |
| MMR 1 = lmer(MMR ~ Temperature*pH + (1 \| tank) + (1\|respirometer), data=mr) | | 0.136 |
| MMR 2 = lmer(MMR ~ Temperature + pH + (1 \| tank) + (1\|respirometer), data=data) | | |
| AS 1 = lmer(AS ~ Temperature*pH + (1 \| tank) + (1\|respirometer), data=data) | | 0.196 |
| AS 2 = lmer(AS ~ Temperature + pH + (1 \| tank) + (1\|respirometer), data=data) | | |
| $O_2$crit 1 = lmer($O_2$crit ~ Temperature*pH + (1 \| tank) + (1\|respirometer), data=data) | | 0.894 |
| $O_2$crit 2 = lmer($O_2$crit ~ Temperature + pH + (1 \| tank) + (1\|respirometer), data=data) | | |

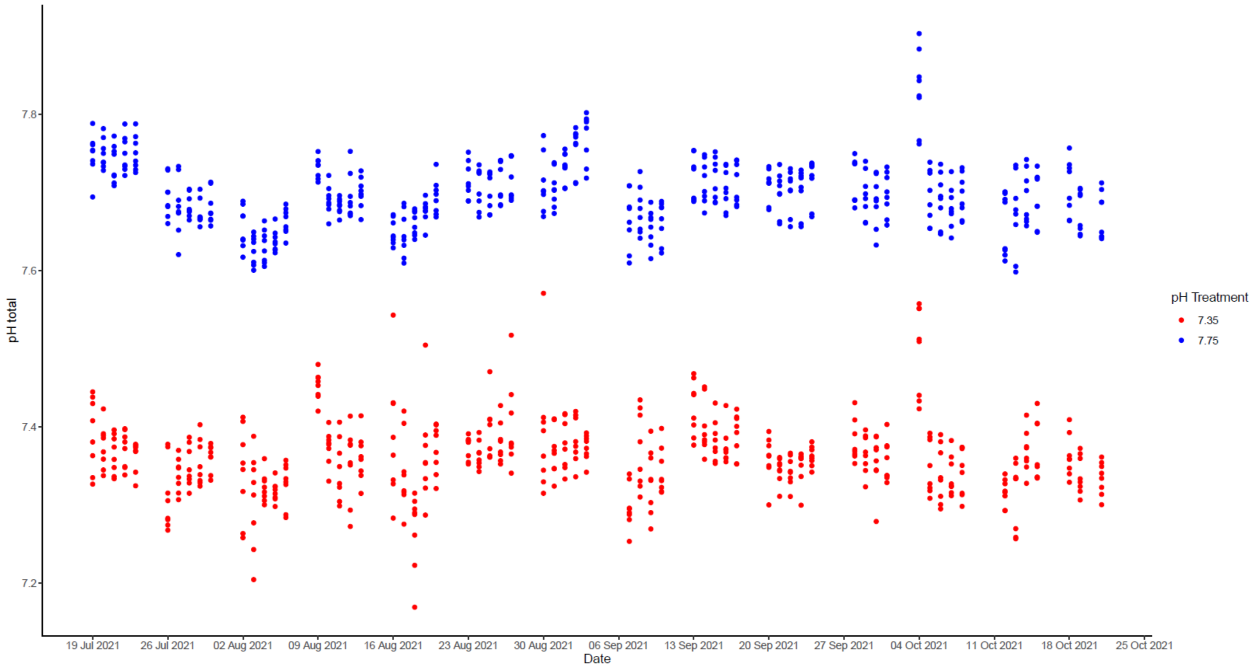

**Figure S1. Values of pH total recorded from Monday to Friday in each of the experimental tanks during the growth period.** In blue: tanks that were kept at a control pH treatment of 7.75. In red: tanks that were kept at an acidified pH treatment of 7.35.

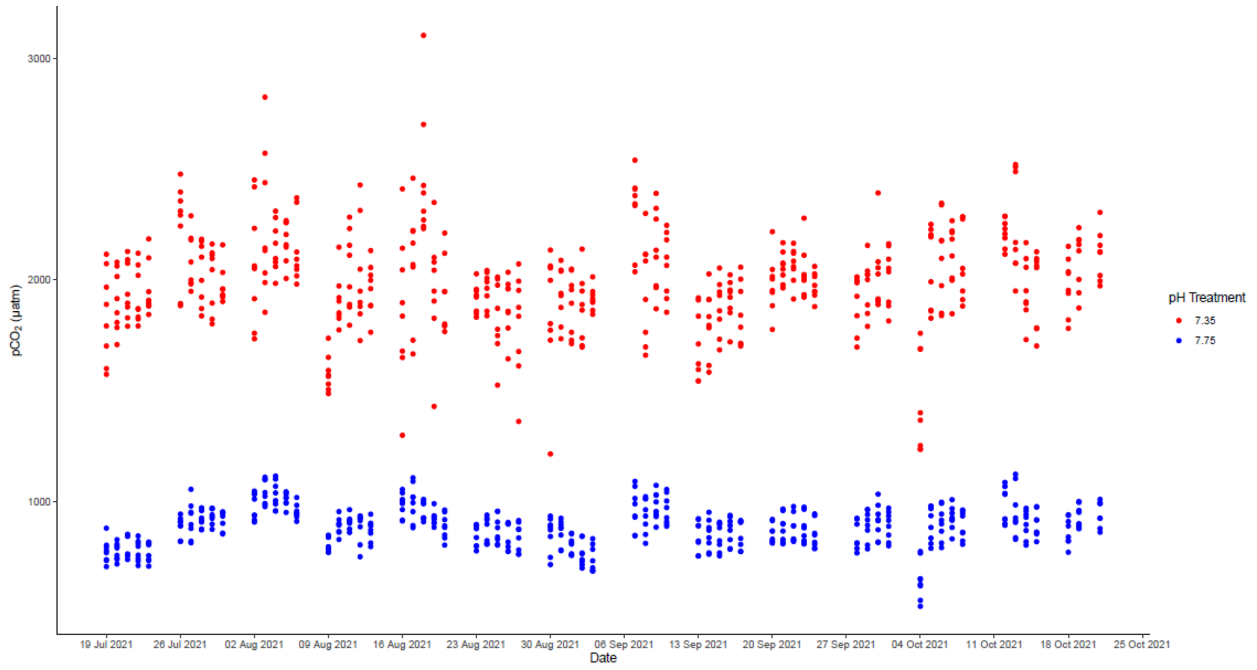

**Figure S2. $pCO_2$ values estimated from pH total measurement and alkalinity for the period where growth was estimated for this experiment.** In blue: tanks that were kept at a control pH treatment of 7.75. In red: tanks that were kept at an acidified pH treatment of 7.35**.**

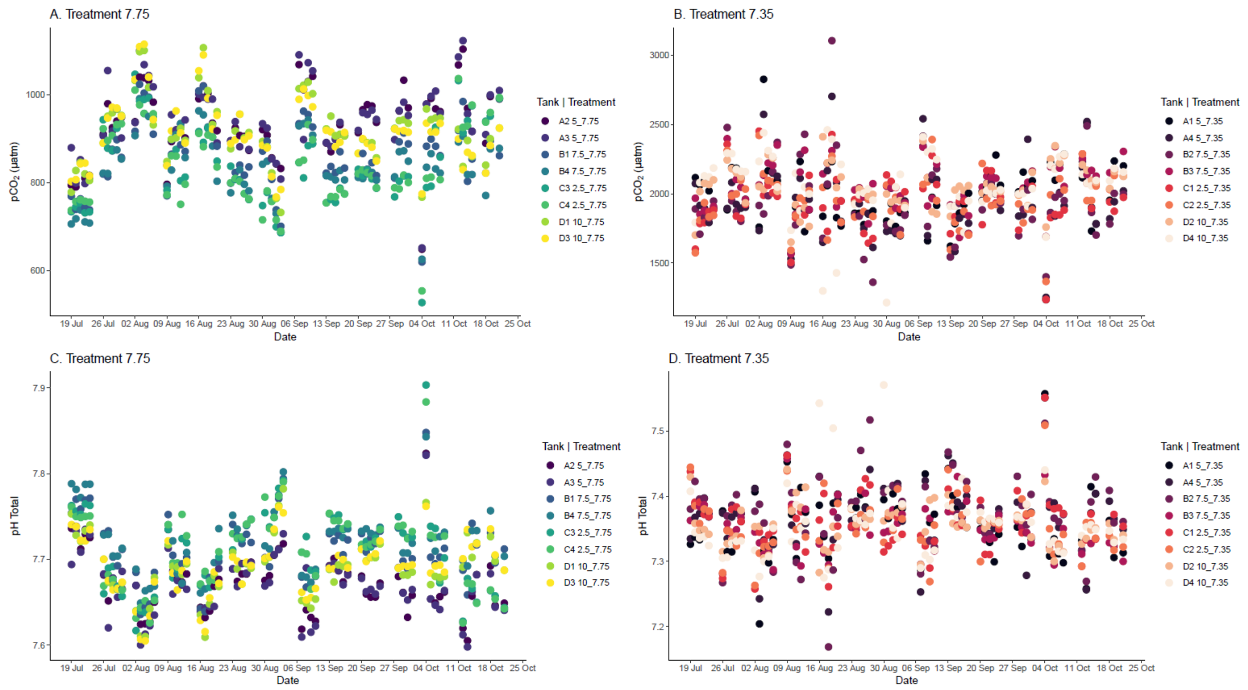

**Figure S3. Seawater monitoring of the experimental tanks throughout the growth period for this experiment. A.** and **B.** $pCO_2$ values estimated from pH total measurement and alkalinity for the period where growth was estimated for this experiment. **A.** tanks that were kept at a control pH treatment of 7.75. **B.** tanks that were kept at an acidified pH treatment of 7.35. **C.** and **D.** Values of pH total recorded everyday from Monday to Friday in each of the experimental tank during the growth period. **C.** tanks that were kept at a control pH treatment of 7.75. **D.** tanks that were kept at an acidified pH treatment of 7.35.

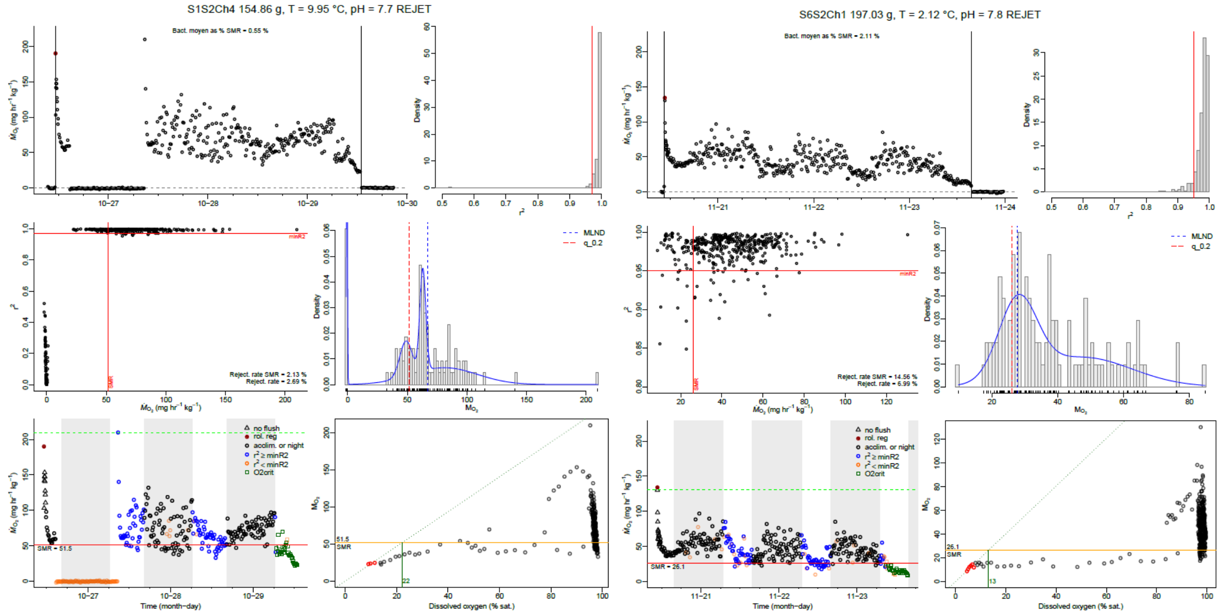

**Figure S4. Evolution of oxygen uptake during 4-day experiments for 2 redfish. A fish which reached stable values typical of SMR during the latter part of the experiment; two fish did not reach SMR.**

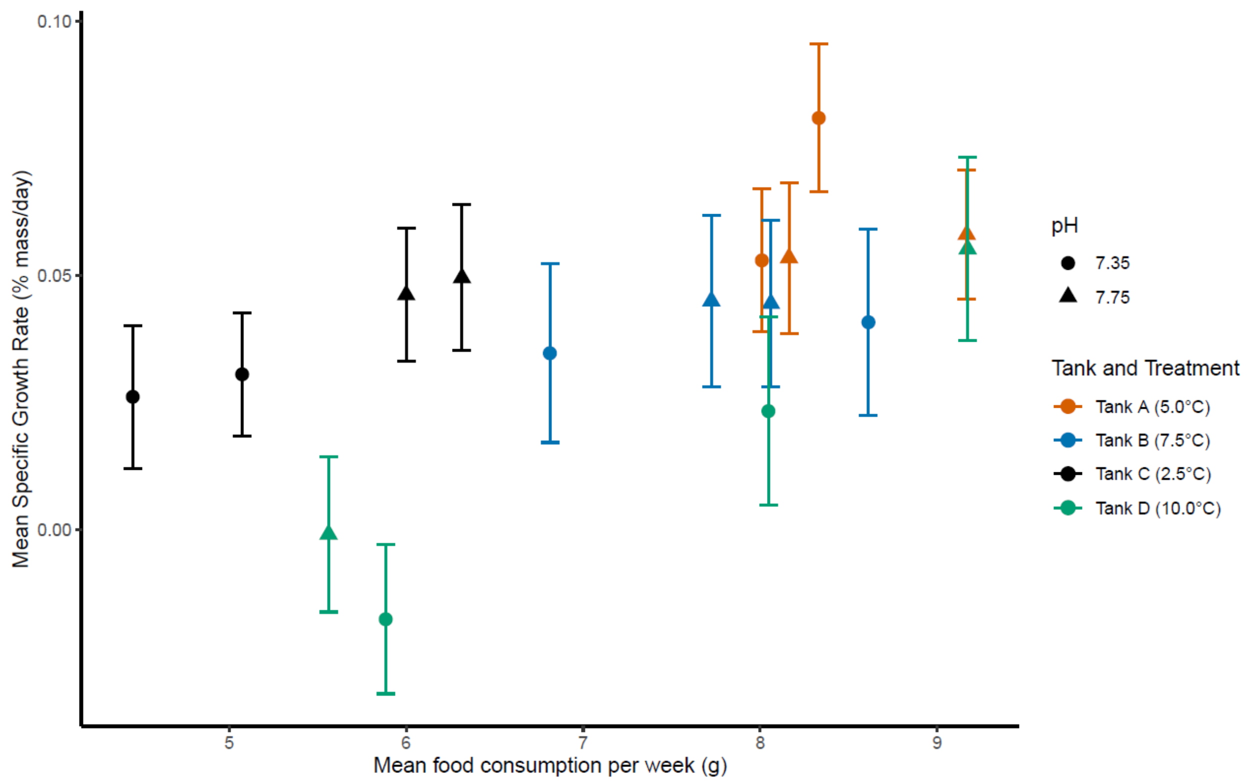

**Figure S5. Mean specific growth rates in function of mean food consumption per week for all tanks. Circles are for the tank that had an acidic pH treatment of 7.35, triangles are for the tanks that had a control pH of 7.75. Each colors represents the 4 different temperature lines (A, B, C and D).**

**Table S5. Mortalities and maximum number of individuals per tanks for the growth experiment from February 22nd to July 13th.**

| Tank | Temperature | pH | Treatment | Max.nbr | Mort |
|---|---|---|---|---|---|
| A1 | 5 | 7.35 | 5_7.35 | 25 | 0 |
| A2 | 5 | 7.75 | 5_7.75 | 25 | 0 |
| A3 | 5 | 7.75 | 5_7.75 | 25 | 0 |
| A4 | 5 | 7.35 | 5_7.35 | 25 | 0 |
| B1 | 7.5 | 7.75 | 7.5_7.75 | 25 | 1 |
| B2 | 7.5 | 7.35 | 7.5_7.35 | 25 | 1 |
| B3 | 7.5 | 7.35 | 7.5_7.35 | 25 | 1 |
| B4 | 7.5 | 7.75 | 7.5_7.75 | 25 | 1 |
| C1 | 2.5 | 7.35 | 2.5_7.35 | 25 | 1 |
| C2 | 2.5 | 7.35 | 2.5_7.35 | 25 | 1 |
| C3 | 2.5 | 7.75 | 2.5_7.75 | 25 | 0 |
| C4 | 2.5 | 7.75 | 2.5_7.75 | 25 | 0 |
| D1 | 10 | 7.75 | 10_7.75 | 25 | 0 |
| D2 | 10 | 7.35 | 10_7.35 | 25 | 2 |
| D3 | 10 | 7.35 | 10_7.35 | 25 | 2 |
| D4 | 10 | 7.75 | 10_7.75 | 25 | 3 |

**Table S6. Mortalities and maximum number of individuals per tanks for the growth experiment from July 18th to October 20th.**

| Tank | Temperature | pH | Treatment | Max.nbr | Mort |
|---|---|---|---|---|---|
| A1 | 5 | 7.35 | 5_7.35 | 42 | 0 |
| A2 | 5 | 7.75 | 5_7.75 | 42 | 0 |
| A3 | 5 | 7.75 | 5_7.75 | 42 | 0 |
| A4 | 5 | 7.35 | 5_7.35 | 42 | 0 |
| B1 | 7.5 | 7.75 | 7.5_7.75 | 42 | 1 |
| B2 | 7.5 | 7.35 | 7.5_7.35 | 42 | 1 |
| B3 | 7.5 | 7.35 | 7.5_7.35 | 42 | 1 |
| B4 | 7.5 | 7.75 | 7.5_7.75 | 42 | 1 |
| C1 | 2.5 | 7.35 | 2.5_7.35 | 42 | 1 |
| C2 | 2.5 | 7.35 | 2.5_7.35 | 42 | 1 |
| C3 | 2.5 | 7.75 | 2.5_7.75 | 42 | 0 |
| C4 | 2.5 | 7.75 | 2.5_7.75 | 42 | 0 |
| D1 | 10 | 7.75 | 10_7.75 | 42 | 0 |
| D2 | 10 | 7.35 | 10_7.35 | 40 | 1 |
| D3 | 10 | 7.35 | 10_7.35 | 42 | 2 |
| D4 | 10 | 7.75 | 10_7.75 | 42 | 4 |
